# Convergent and clonotype-enriched mutations in the light chain drive affinity maturation of a public antibody

**DOI:** 10.1101/2025.03.07.642041

**Authors:** Vishal Rao, Iden Sapse, Hallie Cohn, Duck-Kyun Yoo, Pei Tong, Jordan Clark, Bailey Bozarth, Yuexing Chen, Komal Srivastava, Gagandeep Singh, Florian Krammer, Viviana Simon, Duane Wesemann, Goran Bajic, Camila H. Coelho

## Abstract

Public antibodies that recognize conserved epitopes are critical for vaccine development, and identifying somatic hypermutations (SHMs) that enhance antigen affinity in these public responses is key to guiding vaccine design for better protection. We propose that affinity-enhancing SHMs are selectively enriched in public antibody clonotypes, surpassing the background frequency seen in antibodies carrying the same V genes, but with different epitope specificities. Employing a human *IGHV4-59*/*IGKV3-20* public antibody as a model, we compare SHM signatures in antibodies also using these V genes, but recognizing other epitopes. Critically, this comparison identified clonotype-enriched mutations in the light chain. Our analyses also show that these SHMs, in combination, enhance binding to a previously uncharacterized viral epitope, with antibody responses to it increasing after multiple vaccinations. Our findings offer a framework for identifying affinity-enhancing SHMs in public antibodies based on convergence and clonotype-enrichment and can help guide vaccine design aimed to elicit public antibodies.

**Graphical Abstract:** 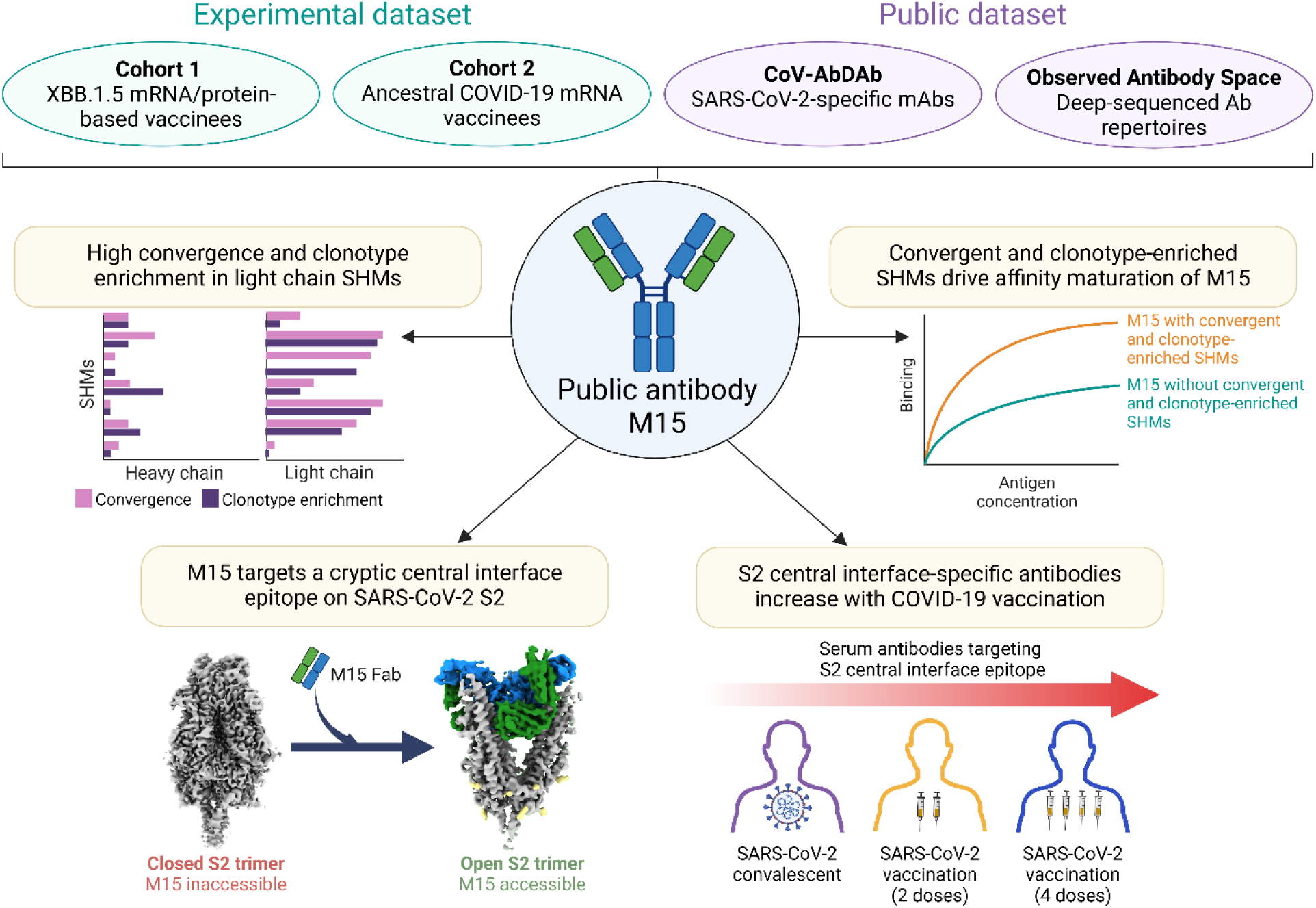

**Highlights:**

- Clonotype-enriched SHMs are identified in the light chain of a public antibody, M15
- These light chain SHMs enhance affinity of M15
- M15 targets a previously undescribed, conserved viral epitope
- Serum antibody levels targeting this epitope increase after repeated vaccinations

## INTRODUCTION

The adaptive humoral immune system recognizes a wide range of antigens due to the vast diversity of the antibody repertoire. Stochastic recombination of germline variable (V), diversity (D), and joining (J) antibody genes give rise to an estimated 10^12^ possible combinations, referred to as clonotypes^1,2^. Despite this extensive diversity, different individuals can generate the same antibodies in response to a given antigen, known as public antibodies. Public antibodies have been identified in response to various pathogens, including severe acute respiratory syndrome coronavirus 2 (SARS-CoV-2), human immunodeficiency virus 1 (HIV-1), influenza virus, dengue virus, Zika virus, and Ebolavirus^3–9^. The recurring presence of a public antibody suggests that certain germline gene segments possess paratope features that improve binding to the antigenic target^10^. In addition to V(D)J recombination, another key feature that contributes to the diversity of antibodies is somatic hypermutation (SHM), which improves affinity for a specific antigen^11–14^. Previous studies have shown that key SHMs acquired during affinity maturation of public antibodies are convergently selected in different individuals^3,15^. Thus, public antibodies serve as a valuable model to investigate how SHM shapes antibody evolution.

Affinity maturation occurs in germinal centers within secondary lymph nodes, where B cells acquire SHMs in the dark zone and then shuttle to the light zone, where selection based on affinity takes place^12,16^. This spatial separation suggests that SHMs in the dark zone are initially acquired independently of their potential to enhance affinity. As a result, SHMs can be neutral, disruptive, or beneficial for affinity maturation, and only a subset of these mutations is retained during clonal expansion. Identifying affinity-enhancing SHMs in the backdrop of other SHMs remains a challenging question.

While affinity-enhancing SHMs are known to be convergently selected across individuals^3,15^, not all convergent SHMs are necessarily affinity-enhancing. Some may be occurring frequently due to the selective targeting of the activation-induced cytidine deaminase (AID) enzyme responsible for SHM, at certain hotspot motifs^17,18^. If these convergent SHMs do not selectively contribute to affinity maturation of a particular clonotype, they would likely appear at a background frequency in B cells, regardless of epitope specificity. This is further supported by the fact that antibodies recognizing different epitopes often utilize the same antibody V gene which contributes to major interactions with epitopes^19^.

In this context, we hypothesize that affinity-enhancing SHMs in public antibody lineages are not only convergent, but also enriched in the public antibody clonotype, compared to clonotypes with other epitope specificities, using the same V gene. We use M15, a public antibody targeting the S2 domain of the SARS-CoV-2 spike protein, as a model to address this hypothesis. Our findings reveal that the affinity maturation of M15-like antibodies is driven by convergent and clonotype-enriched SHMs, which enhance binding to a previously uncharacterized central interface epitope exposed in the open conformation of S2. We also show that serum antibodies targeting this epitopic region increase in proportion upon multiple vaccinations. Overall, our analysis provides a framework for identifying affinity-enhancing SHMs, which can be leveraged to optimize the design of vaccine immunogens that induce favourable affinity maturation pathways.

## RESULTS

### M15 is a human public antibody elicited specifically upon exposure to SARS-CoV-2 in multiple cohorts

The antibody repertoires of 603 single-cell plasmablast sequences of five individuals who received either the XBB.1.5 monovalent mRNA booster (BioNTech/Pfizer or Moderna) or the protein-based booster (Novavax) in 2023 (Cohort 1)^20^ were analyzed to identify public antibodies binding to the spike protein of SARS-CoV-2 (**Table S1**). To identify the most public antibody in Cohort 1, we mapped each sequence from our repertoire to a database of SARS-CoV-2 antibody sequences, named CoV-AbDab^21^ based on identical V and J gene usage and greater than 70% CDR3 amino acid similarity in both heavy and light chains from the repertoire, as this would ensure that the identified public antibodies from different individuals would have evolved from highly similar germline B cells^22,23^. Based on these criteria, four sequences from Cohort 1 (antibodies M13, M14, M15, and M30) mapped to at least one antibody in the database, while none of the antibodies from the protein-based cohort mapped to sequences deposited in CoV-AbDab (**Table S2**). One of the antibodies, M15, was found to map to 17 distinct antibodies reported across seven studies of SARS-CoV-2-infected or vaccinated individuals, suggesting it may be a widely elicited public antibody.

Next, to validate M15 as a public antibody, we screened single-cell V(D)J sequences from plasmablasts and spike-specific memory B cells in a cohort of SARS-CoV-2 mRNA vaccinees (Cohort 2, n = 19) (**Table S1**) and two additional datasets: (i) Observed Antibody Space (OAS), comprising antibody repertoires elicited under different conditions, including tonsillitis, obstructive sleep apnea, HIV-1 infection, and SARS-CoV-2 infection^24^, and (ii) a dataset we compiled in-house, containing 3734 sequences of monoclonal antibodies (mAbs) elicited upon influenza virus infection or vaccination from 24 studies and *Plasmodium falciparum* (malaria) vaccination from two studies (**Table S3**). These sequences were also mapped to M15 based on identical V and J germline gene usage, along with a CDR3 amino acid similarity of 70% or greater in both the heavy and light chains. Only individuals exposed to SARS-CoV-2 through infection or vaccination exhibited M15-like sequences in their repertoire, further confirming the antigen specificity of M15 to SARS-CoV-2 (**Table 1**).

**Table 1.** Sources of M15-like antibody sequences. The database or cohort source for each condition is mentioned in parentheses. Sequences from influenza virus infection/vaccination and malaria vaccination were compiled in-house from 26 independent studies (described in Table S3). The instances where M15-like sequences were identified are highlighted in gray.

| Condition (Cohort) | Total number of sequences | M15-like sequences |
| --- | --- | --- |
| XBB.1.5 mRNA/protein-based vaccination (Cohort 1) | 603 | 1 |
| CoV-AbDab | 11,120 | 17 |
| COVID-19 mRNA vaccination (ancestral strain) (Cohort 2) | 1,794 | 7 |
| SARS-CoV-2 infection (OAS) | 774,147 | 122 |
| Healthy (OAS) | 568,139 | 0 |
| Obstructive sleep apnea (OAS) | 3,327 | 0 |
| Tonsillitis (OAS) | 17,611 | 0 |
| Tonsillitis/Obstructive sleep apnea (OAS) | 10,133 | 0 |
| HIV-1 (OAS) | 5,547 | 0 |
| Multiple sclerosis (OAS) | 179,468 | 0 |
| Influenza virus infection/vaccination | 3,734 | 0 |
| Malaria vaccination | 193 | 0 |

Among four studies involving individuals infected with SARS-CoV-2, all participants in two of the studies^25,26^ harbored M15-like sequences (**Figure S1A**), contributing a total of 122 sequences similar to M15. By combining all M15-like sequences from Cohort 1, Cohort 2, and the CoV-AbDab and OAS databases, we identified a total of 147 M15-like sequences. These sequences were found across various B cell subtypes, including plasmablasts, memory B cells, and naïve B cells in peripheral blood, as well as germinal center B cells and plasma cells in draining lymph nodes (**Figure S1B**).

### M15 binds broadly to the S2 domain of spike protein of sarbecoviruses

We evaluated the binding of M15 to SARS-CoV-2 spike proteins of WA1/2020, XBB.1.5, and JN.1 variants, in addition to human coronaviruses (HCoVs) 229E, HKU1, OC43, NL-63, MERS-CoV, and SARS-CoV (**Figure 1A**). M15 bound to SARS-CoV-1 and all SARS-CoV-2 variants tested but not the other HCoV spike proteins, suggesting broad binding across sarbecoviruses (**Figure 1B**). We then tested the binding of M15 to individual subunits of the SARS-CoV-2 USA-WA1/2020 spike protein and found that it specifically binds to the highly conserved S2 domain (**Figure 1C**). This finding potentially explains its broad binding activity against sarbecoviruses (**Figure 1B**). However, M15 did not neutralize WA1/2020, XBB.1.5, or JN.1 variants *in vitro* (**Figure 1D, S2**) and was unable to protect transgenic mice expressing the human ACE2 receptor (hACE2-K18) against viral challenge (**Figure 1E-G**). We also confirmed that other M15-like antibodies from the CoV-AbDab database bind to the S2 subunit of spike protein of sarbecoviruses and lack neutralization capacity against SARS-CoV-2 (**Figure S2**). Now that we know the antigen that M15 binds to, we addressed our hypothesis that convergent and clonotype-enriched SHMs enhance the affinity of M15 with its cognate antigen, S2.

**Figure 1.**
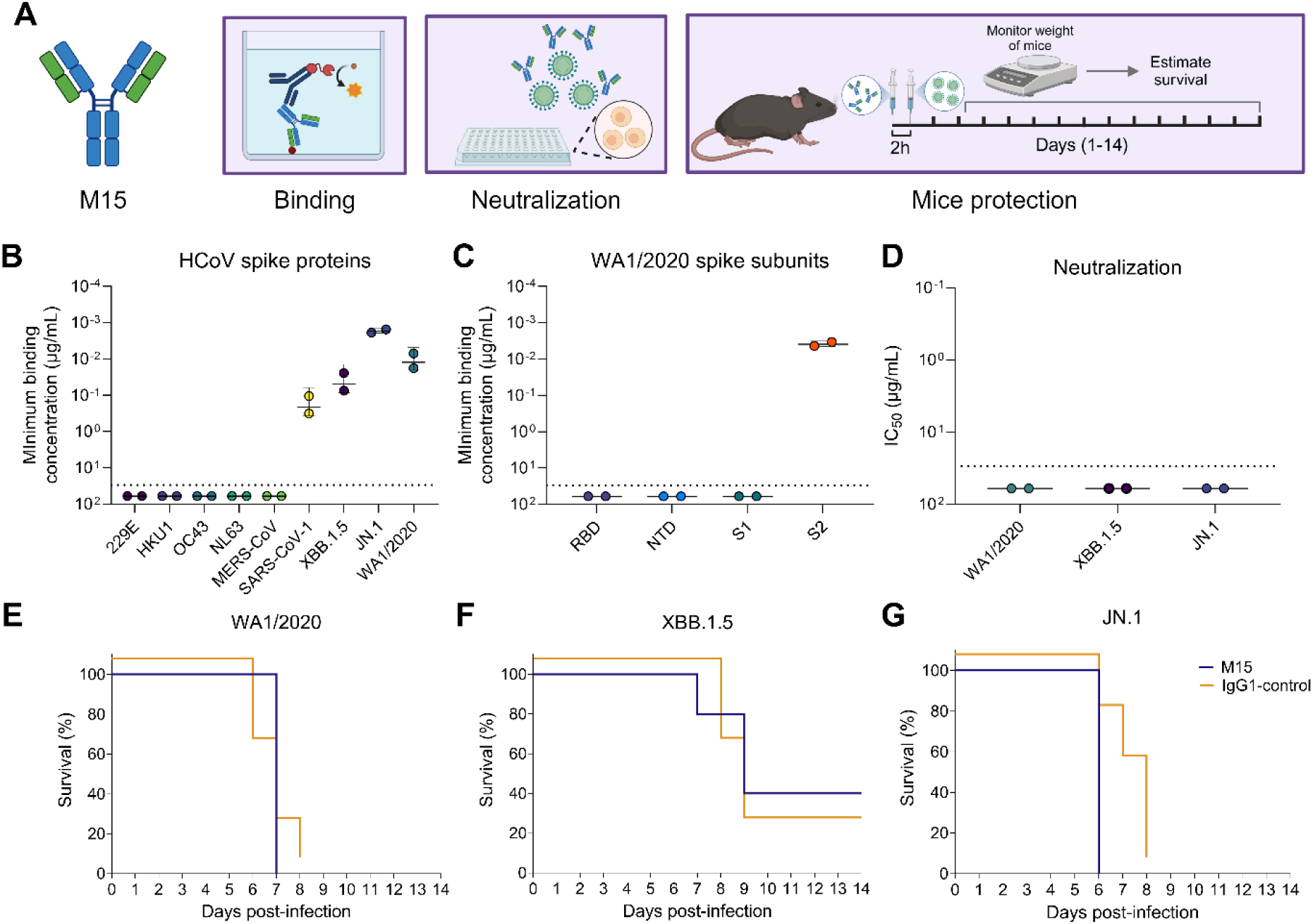
Binding, neutralization, and *in vivo* protection activity of M15. **A**) Schematic of binding, *in vitro* neutralization, and *in vivo* protection experiments performed using M15, created with BioRender. **(B-C)** Binding activity of M15 to (B) spike proteins from human coronaviruses, including the XBB.1.5 and JN.1 variants of SARS-CoV-2, and (C) subunits of WA1/2020 spike protein, represented as the minimum binding concentration (µg/mL) measured by ELISA. The dotted line represents the limit of detection (LOD), set at the starting dilution of 30 µg/mL. All values with a minimum binding concentration of > 30 µg/mL, were set to 60 µg/mL for graphing purposes. **(D)** *In vitro* neutralization capacity of M15 against WA1/2020, XBB.1.5, and JN.1 variants, represented as half-maximal inhibitory concentrations (IC50). All values with an IC50 of > 30 µg/mL, were set to 60 µg/mL for graphing purposes **(E-G)** Survival curves of hACE2-K18 mice prophylactically treated with 10 mg/kg (intraperitoneal) of M15 antibody prior to challenge with a 3xLD50 dose of (E) WA1/2020, (F) XBB.1.5, or (G) JN.1. CR9114, an isotype-matched influenza virus anti-hemagglutinin mAb, was used as a negative control mAb.

### Convergent and clonotype-enriched SHMs are exclusively found in the M15 light chain and drive affinity maturation

We first investigated whether the 147 M15-like sequences—identified by matching M15 with Cohort 2, CoV-AbDab, and OAS—could be leveraged to identify convergent SHMs in this public antibody clonotype. We re-constructed clonal lineages from seven individuals who each presented at least three M15-like sequences, namely Donor-1, Donor-2, SCoV-1, SCoV-10, SCoV-11, and SCoV-13 from OAS and one individual, 368.20.B, from CoV-AbDab. To assess convergence, we shortlisted SHMs observed in at least three of the seven individuals and computed the frequency of these mutations within each lineage (**Figure 2A, B**). This yielded five and eight convergent SHMs in the heavy and light chains, respectively. Most of the convergent mutations were contributed by Donor-1, Donor-2, and 368.20.B, which could be explained by higher overall SHM in these individuals (**Figure S3A, B**).

**Figure 2.**
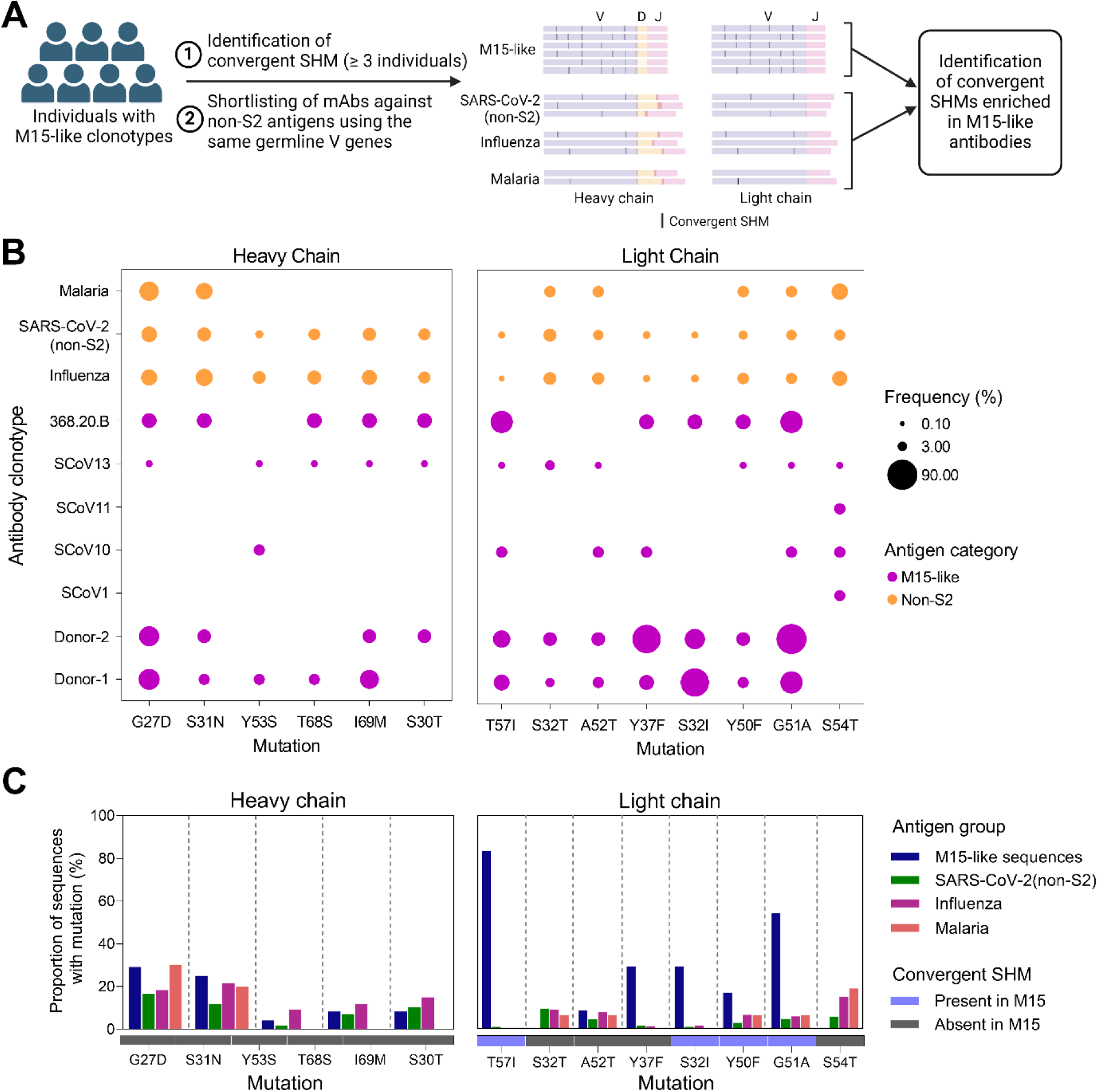
Convergent and specific SHM in M15-like antibodies. **(A)** Schematic workflow for identifying SHMs that are convergent and specific to M15-like antibodies, created with BioRender. Black traces refer to convergent SHMs. The number of sequences illustrated per group is not representative of the actual numbers. Since the non-M15-like sequences (SARS-CoV-2 non-S2, influenza virus, and malaria) share the same germline V gene (*IGH4-59* and *IGKV3-20*) but have different D and J genes, the lengths of the D and J gene segments vary in the cartoon. **(B)** Frequencies of heavy (left) and light (right) chain SHM in M15-like clonal lineages (126 sequences) and negative-control malaria (heavy chain: 10 sequences, light chain: 16 sequences), influenza virus (heavy chain: 134 sequences, light chain: 441 sequences), and non-S2 (heavy chain: 208 sequences, light chain: 648 sequences) mAbs from CoV-AbDab using the same germline V genes. Mutations present in at least three individuals are shown. **(C)** The bar plots represent the frequencies of heavy (left) and light (right) chain SHM in M15-like singleton sequences from Cohort 1, Cohort 2, and CoV-AbDab (21 sequences not previously included in panel B), as well as negative-control antibodies against malaria, influenza virus, and non-S2 epitopes from CoV-AbDab, all using the same germline V genes as above. Underneath the X-axis, a heatmap illustrates the presence or absence of the convergent SHM observed in the original M15 sequence.

Next, to assess clonotype-enrichment of these convergent SHMs in the heavy and light chains, we compared the frequencies of the convergent mutations with mAbs using the same heavy (*IGHV4-59*) or light (*IGKV3-20*) chain V gene as M15 respectively, and specific to non-S2 epitopes from three groups: (i) influenza virus infection/vaccination, (ii) malaria-vaccination, and (iii) SARS-CoV-2-specific mAbs from CoV-AbDab recognizing epitopes outside the S2 domain, henceforth referred to as the non-S2 group. These mAbs share the same heavy or light chain V gene as M15, but belong to clonotypes with distinct specificities due to variations in junctional regions and heavy/light chain pairings^27^. Notably, the frequencies of convergent heavy chain mutations showed no significant differences between M15-like clonotypes and mAbs from the non-S2 group. On the other hand, the light chain mutations S32I, Y37F, Y50F, G51A, and T57I were significantly more frequent in M15-like clonotypes (**Figure 2B**). We also compared the frequencies of these convergent mutations in heavy and light chains of M15-like sequences from Cohort 1, Cohort 2, and CoV-AbDab with the non-S2 group (**Figure 2C**). Again, in this second analysis, heavy chain mutations showed no differences in frequency between groups, while light chain mutations S32I, Y37F, Y50F, G51A, and T57I were enriched in M15-like antibodies (**Figure 2C**). While none of the convergent mutations in the heavy chain appeared in the original sequence of M15, four out of the eight convergent mutations in the light chain, namely, S32I, Y50F, G51A, and T57I, were present in M15 (**Figure 2C**).

Having identified these four mutations in M15, we tested our hypothesis that these convergent and clonotype-enriched SHMs enhance affinity to the antigen, S2. We individually reverted the three most convergent mutations present in M15—S32I, G51A, and T57I—back to their germline residues in the light chain. These mutations were evaluated as single mutants and in combination (triple mutant) to assess their impact on binding affinity to the S2 region of wild-type SARS-CoV-2 (**Figure 3A-C**). While the single mutants showed a slight decrease in binding affinity (S32I, 1.45-fold; G51A, 1.88-fold; T57I, 1.02-fold) compared to M15 (K_D_ 95% C.I, 246.3 – 709.7 nM), the triple mutant showed a 12-fold decrease, roughly matching the K_D_ of the unmutated common ancestor (UCA; K_D_ 95% C.I, 3560 – 8522 nM), implying a synergistic effect of mutations S32I, G51A, and T57I in affinity maturation of M15 (**Figure 3B, C**).

**Figure 3.**
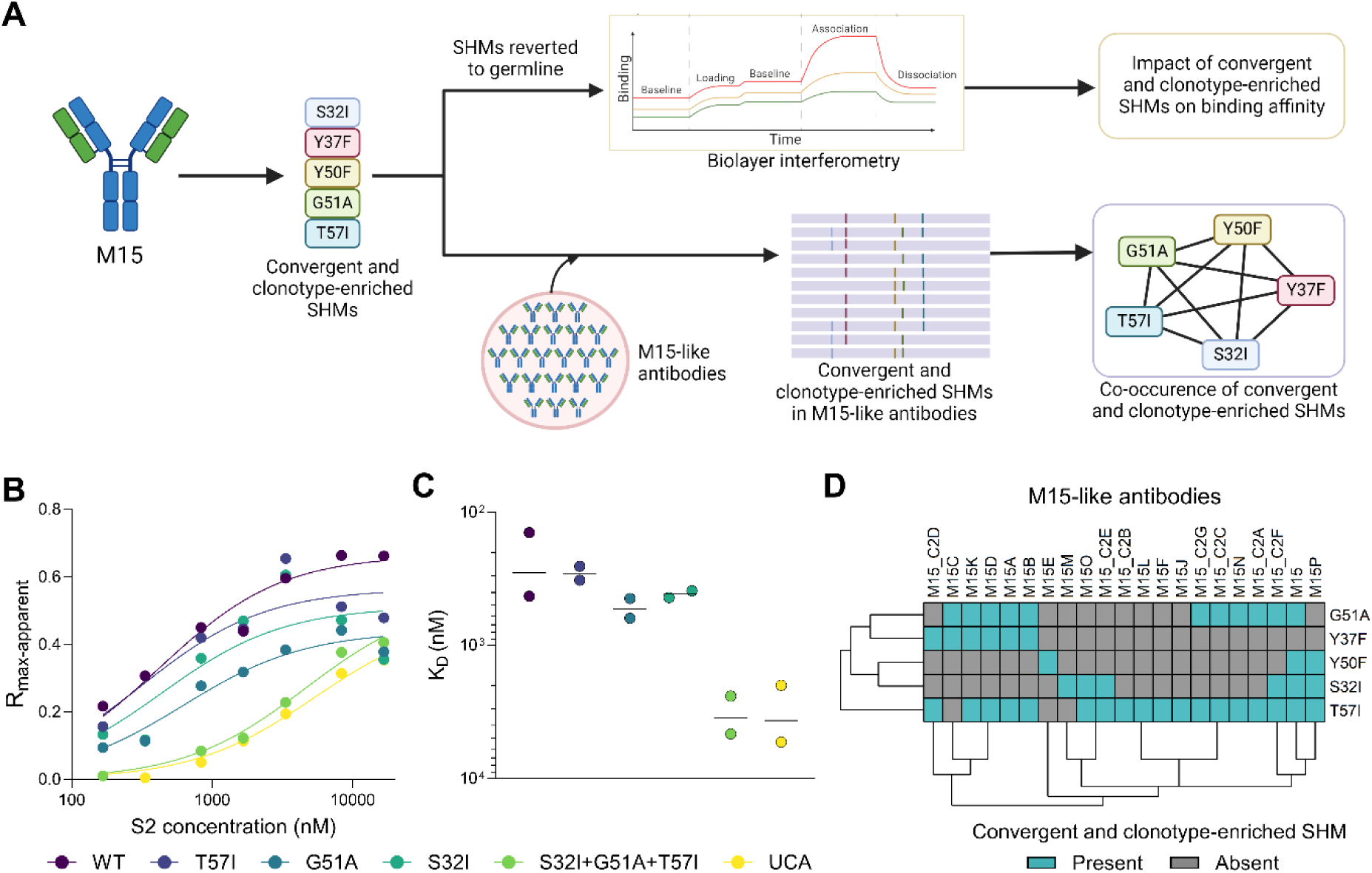
Influence of convergent and clonotype-enriched SHMs on binding affinity of M15. **(A)** Schematic workflow for assessing the role of convergent and clonotype-enriched mutations in affinity maturation of M15 and co-occurrence of these SHMs in M15-like antibodies, created with BioRender. **(B)** Biolayer interferometry (BLI) is used to assess the binding of M15; single reverted mutants G51A, T57I, and S32I; triple mutant with all three mutations reverted; and the unmutated common ancestor (UCA) with S2. Rmax-apparent represents the baseline-corrected maximum binding signal reached during the association step of the assay, measured at different concentrations of S2, ranging from 166.7-16,666.7 nM. The lines represent the best-fit one-site specific binding curves. The BLI assay was performed in duplicates for each antibody (only one of the replicates shown). **(C)** Binding affinity values of M15, the single mutants, the triple mutant, and the UCA, computed as the dissociation constant, KD. KD values were obtained from the best-fit one-site specific binding in GraphPad Prism independently for the duplicates. **(D)** Biclustered heatmap representing the occurrence of convergent and clonotype-enriched SHMs in M15-like antibodies from Cohort 1 (M15), Cohort 2 (M15_C2A-F), and CoV-AbDab (M15A-Q). Biclustering was performed using k-means clustering in the ‘pheatmap’ package in R.

To explore this further, we examined whether convergent and clonotype-enriched SHMs are acquired simultaneously during affinity maturation. None of the M15-like antibodies from Cohort 2 and CoV-AbDab presented all five convergent and clonotype-enriched mutations (**Figure 3D**). Among the M15-like antibodies, the original M15 antibody from Cohort 1 presented the highest number of convergent and clonotype-enriched SHMs (n = 4) (**Figure 3D**). We then conducted a linkage analysis to examine the co-occurrence of convergent mutations in both heavy and light chains. Specifically, for each pair of convergent mutations, we created a contingency table to assess their presence within individual lineages. We computed an odds ratio (O.R.) of co-occurrence for each pair of mutations and considered two mutations to be linked in a lineage if the O.R. was greater than 2 (**Figure S4A-F**). Among the pairs of convergent mutations, G51A-T57I, which was a pair convergent and enriched in M15-like antibodies, exhibited the highest linkage (present in five lineages), followed by G51A-A52T (present in four lineages) (**Figure S4F**). These results indicate that convergent and clonotype-enriched SHMs are critical in the process of affinity maturation of M15. Given that the convergent and clonotype-enriched SHMs enhanced affinity of M15, we hypothesized that these mutations facilitate critical contacts between M15 and S2 from SARS-CoV-2.

To address this, we obtained a 3.4 Å resolution cryogenic electron microscopy (cryo-EM) reconstruction of the M15 Fab bound to a stabilized prefusion SARS-CoV-2 spike S2 trimer^28^. The M15-S2 complex presented stoichiometric threefold symmetry, with one Fab unit bound to each protomer of the S2 trimer (**Figure 4A**). The structure disclosed a previously uncharacterized epitope situated on and around a segment of the S2 central helix (CH) (**Figure 4B**). This epitope, which we refer to as the ‘central interface (CI) epitope,’ is noteworthy for its location at a site of inter-protomer contacts in the canonical ‘closed’ structure of the prefusion S2 homotrimer (**Figure S5A**). The CI epitope is accordingly occluded in the closed prefusion spike conformation, wherein central alpha helices assemble to form a triple-helix coiled-coil. The epitope is thus only accessible in the open spike conformation^29^, which differs markedly from the piriform S2 trimer observed in both the closed full-length spike complex and the unliganded stabilized prefusion S2 trimer. As the CI epitope is only accessible on the open S2, interaction between M15 and S2 favors the open S2 trimer, splaying protomers centrifugally outward from the threefold axis of the trimer (**Figure S6**). In agreement with our binding data demonstrating that M15 reactivity is limited to sarbecoviruses (**Figure 1B**), the contact residues on S2 are well conserved across sarbecoviruses, but divergent across more distantly related HCoVs (**Figure S5C, D**).

**Figure 4.**
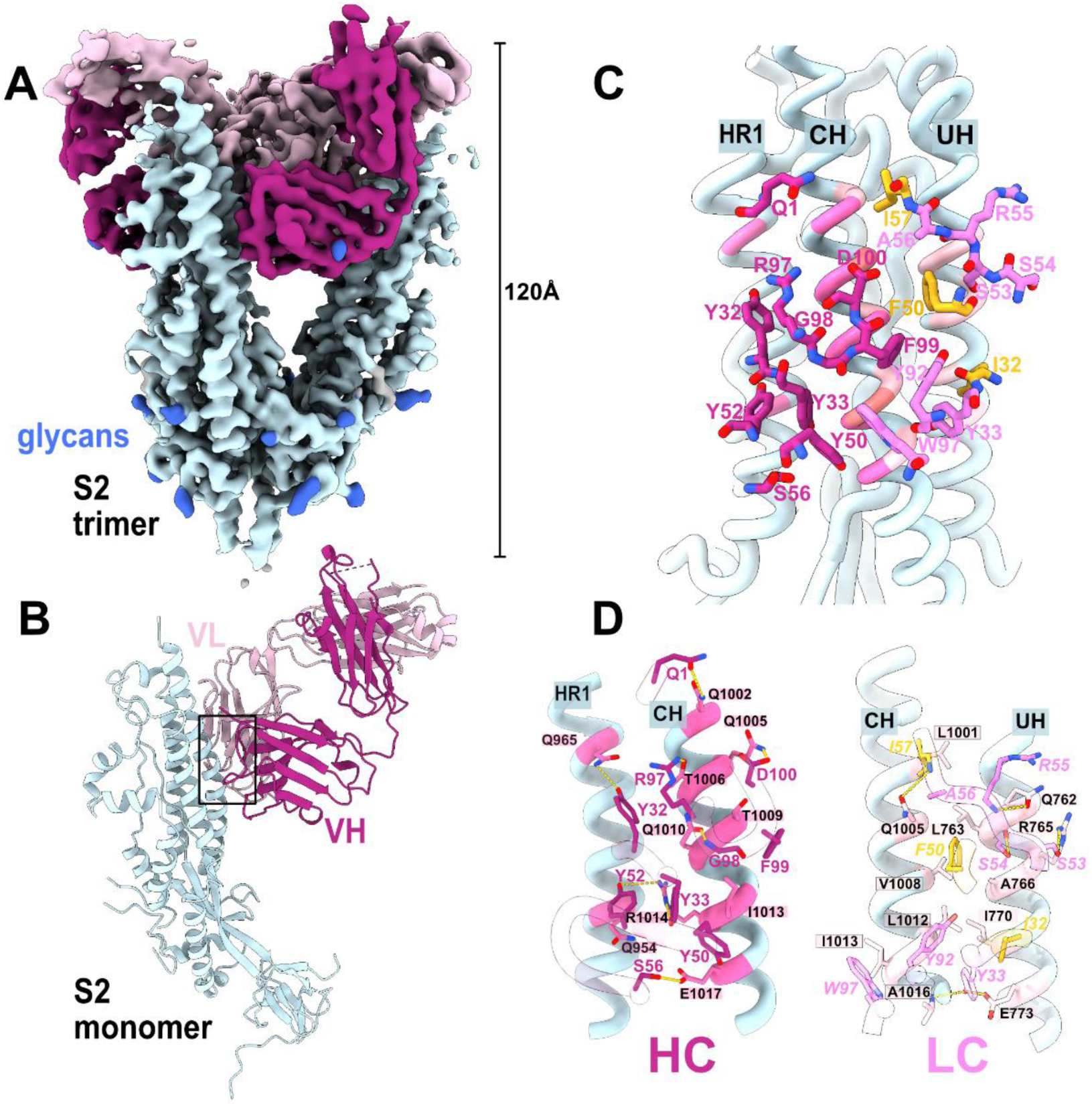
Cryo-EM structure of M15 in complex with S2. **(A)** Density map of the trimeric S2-M15 Fab complex, highlighting S2 (light blue), M15 heavy chain (magenta), M15 light chain (pink). Glycan densities are displayed in royal blue. **(B)** Ribbon representation of S2 protomer-M15 Fab complex structure. **(C)** S2-M15 interface overview (region designated within black box in panel B. S2 Helices (ribbons) are labeled to designate relative locations of heptad repeat 1 (HR1), central helix (CH), and upstream helix (UH). S2 residues are colored according to whether they form contacts with correspondingly colored heavy chain (magenta) or light chain (pink). S2 residues contacting both heavy and light chains are colored coral. Convergent and clonotype-enriched mutations are colored yellow. **(D)** Close-up view of M15 heavy chain (HC; left) and light chain (LC; right) binding interfaces with S2. S2 residues are colored, as in panel C, according to whether they form contacts with correspondingly colored heavy chain (magenta) or light chain (pink). Convergent and clonotype-enriched mutations are colored yellow.

We then analyzed the contacts of M15 Fab with the CI epitope, to address our hypothesis that convergent and clonotype-enriched SHMs facilitate critical contacts with S2. The bound M15 Fab occupies 1315 Å^2^ of buried surface area, with contacts spread across CH and nearby residues on two other alpha helices belonging to heptad repeat 1 (HR1) and the ‘upstream helix (UH)’ (upstream of fusion peptide^30^) (**Figure 4C, D**). Contacts were observed between both heavy and light Fab chains and CH (residues 1001-1016). Three of the convergent and clonotype-enriched SHMs, namely S32I, Y50F and T57I, indeed contributed to critical contacts with the CH and UH (**Figure 2B-C, 4C-D**). These mutations replaced polar residues with hydrophobic ones, favoring thermodynamic stability. The mutated I57 residue in the light chain, which showed the highest convergence and clonotype-enrichment (**Figure 2C**), also formed a ternary interaction with CH residue Q1005, in combination with heavy chain D100 via hydrogen bonds. Thus, the structural data validated that convergent and clonotype-enriched mutations help to drive the interaction between M15 and S2.

Additional contacts, inferred through proximity (<4 Å) and side-chain functional group biochemical complementarity, were observed for heavy chain and light chain with HR1 and UH, respectively. Heavy chain contacts (Q1, Y32, Y33, Y50, Y52, S56, R97, G98, F99, D100) primarily comprise polar interactions with CH (1002-1017). Two contacts (Q1 and Y52) occur between heavy chain and HR1 residues 961 and 965. Light chain contacts (I32, Y33, F50, S53, S54, R55, A56, I57, Y92, G93, W97) are spread between CH (LC Y33, F50, A56, I57, Y92, G93, W97) and UH (I32, Y33, F50, S53, S54, R55). Light chain residues support an abundance of hydrophobic interactions in addition to the convergent and clonotype-enriched SHMs (I57, F50, I32, Y92, W97) and spatial complementarity with a correspondingly hydrophobic cleft formed between CH and UH (S2 residues Y756, F759, L763, A 766, L767, I770, L1001, L1004, V1008, L1012, I1013, A1015, A1016) (**Figure S5B**). An additional ternary interaction occurs at CH residue 1013 via hydrophobic contacts with heavy and light chains (**Figure 4C**). This clasping of CH appears to be enhanced by inter-Fab chain hydrogen bonds (LC Y37-HC F99; LC Q39-HC Q39; **Figure S7**). Now that we have demonstrated that M15, aided by convergent and clonotype-enriched SHMs, binds to a previously uncharacterized epitope on S2, we sought to investigate how antibodies targeting this epitopic region are shaped by varying levels and modes of antigen exposure.

### Serum antibody levels targeting the CI epitope rise with repeated COVID-19 vaccinations

We assessed extent of competition between the M15 mAb and sera of COVID-19 convalescent or vaccinated individuals to estimate and compare the prevalence of antibodies targeting the CI epitope (**Figure 5A**). Human sera samples (Cohort 3) were obtained from four exposure groups: pre-pandemic (n = 24), SARS-CoV-2 convalescent (n = 25), two doses of SARS-CoV-2 vaccination (2X vaccination) (n = 24), and four doses of SARS-CoV-2 vaccination (4X vaccination) (n = 12) (**Table S1, S4**). Among the exposure groups, individuals in the 4X vaccination group exhibited the highest competition with M15, with 91.7% (11 out of 12) classified as responders (median effective dilution (ED_50_) > 1) (**Figure 5B, S8A**). In contrast, only one individual (4.16%) from the pre-pandemic group was identified as a weak responder (**Figure 5B, S8A**). CI epitope competition in the convalescent and 2X vaccination groups was low, with only 12% (3/25) and 16.6% (4/24) responders, respectively (**Figure 5B, S8A**).

**Figure 5.**
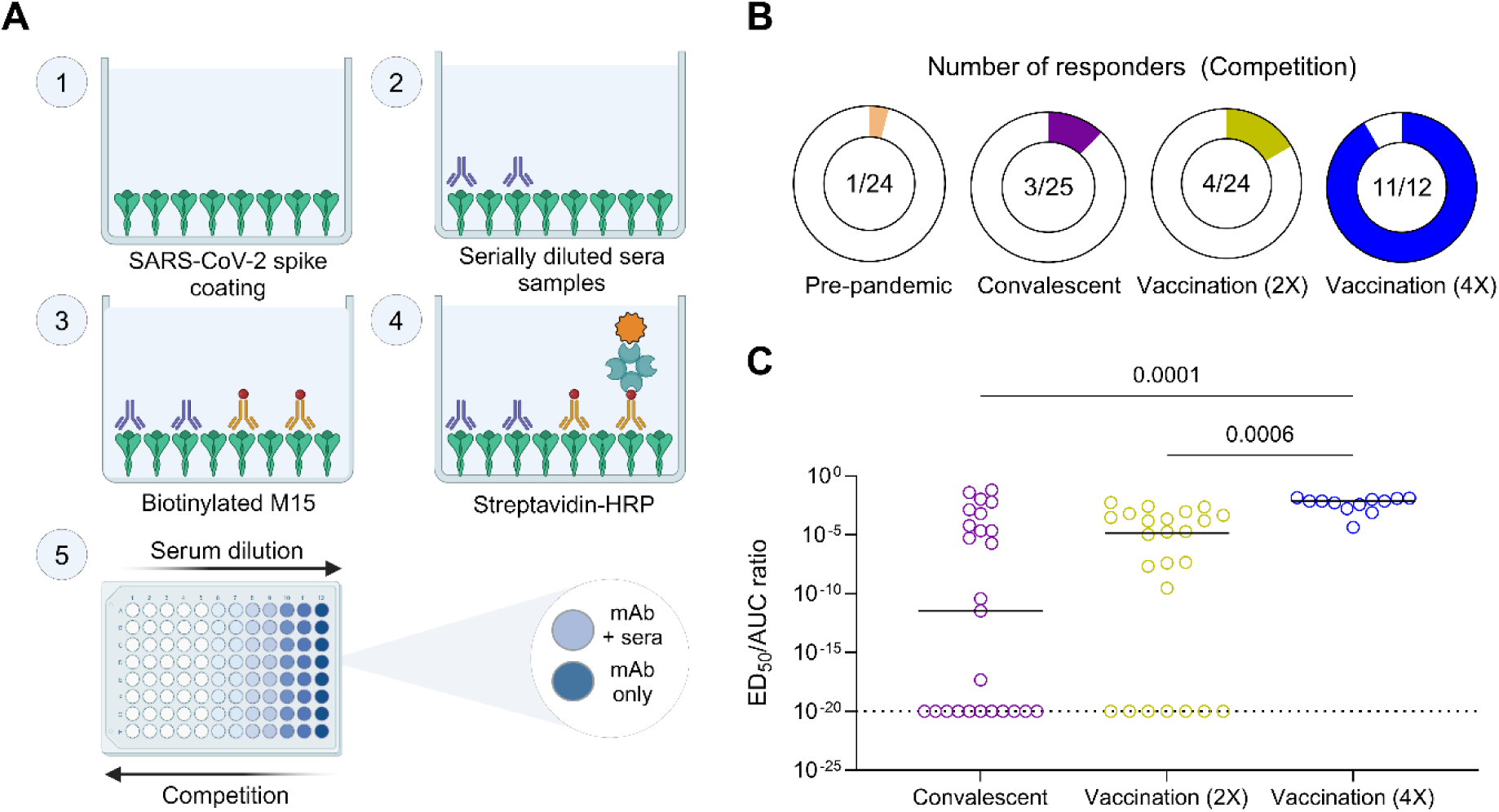
Prevalence of sera antibodies targeting the S2 central interface epitope in SARS-CoV-2-infected and vaccinated individuals. **(A)** Schematic workflow of the protocol employed for the serum competition ELISA, created with BioRender. **(B)** Proportion of responders among the samples tested in a serum competition ELISA between M15 (biotinylated) and human sera from pre-pandemic samples (n = 24) or SARS-CoV-2 convalescent samples (n = 25), two doses of SARS-CoV-2 vaccination (n = 24), or four doses of SARS-CoV-2 vaccination (n = 12). Responders are identified as individuals showing competition ED50 values greater than 1. **(C)** The ratio of competition ED50 to anti-spike serum AUC values are depicted across individuals from the convalescent, vaccination (2X), and vaccination (4X) exposure groups. ED50/AUC values less than 10^-20^ were set to a detection limit of 10^-20^. Comparisons between exposure groups were performed using the Kruskal-Wallis test followed by Dunn’s multiple correction. All serum competition ELISAs and serum ELISAs were performed in duplicates, and the ED50 values and AUCs were calculated as geometric means of the replicates.

We then investigated whether higher serum competition with M15 was a mere consequence of higher anti-spike serum antibody titers, or if the proportion of serum antibodies targeting this epitopic region increases with multiple vaccinations (**Figure S8B**). To answer this, we normalized the extent of competition (i.e., ED_50_) with the serum antibody titers calculated as the area under the curve (AUC). While the extent of competition correlated positively with the serum antibody titers (ED_50_ vs. AUC Spearman correlation coefficient (r_s_) = 0.5586, *p* < 0.001) (**Figure S8C**), the 4X vaccination group still showed significantly higher normalized competition as compared to the 2X vaccination (Kruskal-Wallis test; *p* = 0.0006) and convalescent (*p* = 0.0002) groups (**Figure 5C**). This indicates that the proportion of serum antibodies targeting the CI epitope increases with each vaccine dose.

## DISCUSSION

We hypothesized that affinity-enhancing SHMs would be selectively enriched in a public antibody clonotype beyond the background frequency observed in clonotypes targeting other epitopes using the same V gene. We addressed this hypothesis using a human public antibody model, M15, elicited upon SARS-CoV-2 infection or COVID-19 vaccination, which uses the *IGHV4-59*/*IGHJ4* and *IGKV3-20*/*IGKJ1* heavy and light chain V/J genes, respectively. Notably, previous studies have identified public clonotypes using the same V and J genes and CDR3 sequences as M15 that are also elicited upon SARS-CoV-2 exposure^3,23,31^. Using 147 M15-like sequences, we identified five convergent and clonotype-enriched SHMs exclusively in the light chain. Reversion of three of these SHMs, S32I, T57I, and G51A to their germline residues resulted in a 12-fold decrease in binding affinity, suggesting that these convergent and clonotype-enriched SHMs drive affinity maturation of M15. The minimal change in affinity following individual residue reversions suggests that binding interactions are driven by the synergistic effect of these linked mutations. The role of G51A in the triple revertant loss of affinity remains particularly puzzling, as our cryo-EM structure suggests that A51 is not a direct contact residue. Considered in tandem with the linkage of G51A with T57I, our data point to the importance of analyzing individual contacts within the broader context of epistatic effects on antigen-antibody interactions^32–34^.

Previous studies have demonstrated that primary antibody sequence diversity, embedded in specific motifs, imparts pre-determined specificity to certain antibody germlines, as these motifs are capable of directly binding the epitope in their unmutated state^10,35–37^. To our knowledge, this is the first comparative study that highlights SHM as a potential determinant of antibody specificity by comparing frequencies of SHMs within V genes across antibodies with different specificities. However, it is important to note that this conclusion is applicable within the context of rearranged heavy and light chain antibody germlines—specifically the M15 clonotype in our case. We cannot entirely rule out the possibility that convergent SHMs enriched in M15 might also contribute to affinity maturation in *IGKV3-20* antibodies using other light chain J genes or heavy chain V(D)J genes. Nonetheless, this is unlikely, given the substantial enrichment of these SHMs within the M15 clonotype compared to other *IGKV3-20* antibodies.

Affinity-enhancing SHMs have previously been shown to be convergently selected in public antibodies^3,15^, raising the question of how much the concept of clonotype-enrichment helps narrow down affinity-enhancing SHMs, beyond simple convergence. This is particularly evident when comparing the frequencies of heavy chain SHMs G27D and S31N (**Figure 2C**) to light chain SHMs Y50F and S32I. While all four SHMs exhibit similar levels of convergence, Y50F and S32I show significant clonotype-enrichment. In contrast, G27D and S31N have convergence rates comparable to background frequencies. Structural analysis further supported this distinction, revealing that only Y50F and S32I are involved in contacts, while G27D and S31N are not (**Figure 4C, D**), reinforcing that clonotype-enrichment is a characteristic feature of affinity-enhancing SHMs.

Our structural analysis also revealed that M15 recognizes a previously undiscovered C.I epitope accessible on the open configuration of S2. We observed a vaccine dose-dependent increase in sera antibodies targeting the S2 CI epitope, with individuals who received four COVID-19 vaccine doses showing a higher proportion of these antibodies compared to those who received two doses or those convalescent from COVID-19 infection. Further investigation would be needed to determine if other antibodies (if any) targeting this epitope are also non-protective like M15, and if they increase in proportion upon multiple vaccinations.

In conclusion, our study emphasizes that affinity maturation of public antibodies is driven by convergent and clonotype-enriched SHMs, and our work reveals a previously undescribed CI epitope in S2 of sarbecoviruses. Identifying other public antibodies, not only against viruses but also against other pathogens, could provide valuable mechanistic insights into affinity maturation. This could further inform the design of immunogens that drive favorable affinity maturation pathways and elicit affinity-enhancing SHMs that improve the neutralization potency and protection of antibodies against a broader range of pathogens.

### Limitations of the study

While our study demonstrates that convergent and clonotype-enriched SHMs play a crucial role in affinity maturation, it is currently limited to the M15 clonotype. Further investigation is needed to determine whether this phenomenon extends to other clonotypes. Additionally, although we identified M15-like antibodies following SARS-CoV-2 infection and vaccination with the ChAdOx1 nCoV-19 vaccine (an adenoviral vector-based vaccine), the serological profiles of antibodies targeting the central interface epitope following non-mRNA-based COVID-19 vaccines or breakthrough infections, have yet to be characterized.

## RESOURCE AVAILABILITY

### Lead contact

Further information and requests for resources and reagents should be directed to and will be fulfilled by the lead contact, Camila H. Coelho

### Materials availability

All reagents will be made available upon request after completion of a Materials Transfer Agreement.

### Data and code availability

Heavy and light chain nucleotide sequences from plasmablasts in Cohort 1 and plasmablasts and memory B cells in Cohort 2 have been deposited on the National Center for Biotechnology Information (NCBI) portal (BioProject: PRJNA1134144). The EM maps have been deposited in the Electron Microscopy Data Bank (EMDB) under accession code EMD-48507 and the accompanying atomic coordinates in the Protein Data Bank (PDB) under accession code 9MPW. Code used for the mutation analysis has been deposited on GitHub (https://github.com/coelholab/Rao_et_al_2025.git). Additional data supporting the findings can be found in the supplementary material.

## Acknowledgments

This project was funded by the Icahn School of Medicine at Mount Sinai and partially funded by the National Institutes of Health (U19 AI142737, U19 AI168631 to CHC as IOF Project leader, AI165072 and AI170715 to DRW) and NIH FIRST (U54CA267776 to CHC). We acknowledge support from NIH R01 AI168178 and the Irma T. Hirschl/Monique Weill-Caulier Trust to G.B. Some of this work was performed at the National Center for CryoEM Access and Training (NCCAT) and the Simons Electron Microscopy Center located at the New York Structural Biology Center, supported by the NIH Common Fund Transformative High Resolution Cryo-Electron Microscopy program (U24 GM129539) and by grants from the Simons Foundation (SF349247) and NY State Assembly. This work was supported in part through the computational and data resources and staff expertise provided by Scientific Computing and Data at the Icahn School of Medicine at Mount Sinai and supported by the Clinical and Translational Science Awards (CTSA) grant UL1TR004419 from the National Center for Advancing Translational Sciences. Research reported in this publication was also supported by the Office of Research Infrastructure of the National Institutes of Health under award number S10OD026880 and S10OD030463. We thank the Flow Cytometry Core Facility and staff at the Icahn School of Medicine at Mount Sinai for their assistance. The BSL-3 facility used in this project for work with SARS-CoV-2 virus stocks is a NIH BSL-3/ABSL-3 facility, part of the BSL-3 Biocontainment CoRE. This core is supported by funding from the ISMMS Dean’s Office and investigator contributions through a cost recovery mechanism. The facility use reported in this publication was supported by the National Institute of Allergy and Infectious Diseases of the National Institutes of Health under award number G20AI174733 (R.A. Albrecht). The content is solely the responsibility of the authors and does not necessarily represent the official views of the National Institutes of Health. We thank Dr. Ali Ellebedy at Washington University in St. Louis for generously sharing information on SARS-CoV-2 antibody sequences from germinal centers that were used in this study. We acknowledge Dr. Sachin Kumar, Dr. Ali Sanjari Moghaddam and Dr. Fang Tian from the Wesemann laboratory for their assistance with sample processing for Cohort 2. We thank Dr. Raianna Fantin and Dr. Alana Tiffney from the Coelho laboratory for their valuable feedback on the manuscript.

## Author Contributions

VR, DRW and CHC designed the study. VR, IS, HC, DKY, PT, JC and BB performed experiments. VR, IS and JC analyzed data. DKY, KS, YC, VS and DRW provided access to key biosamples and metadata from participants. GS and FK provided critical materials for this study. GB and CHC supervised the study. DRW, GB and CHC provided funding for the study. VR, IS, GB and CHC wrote the manuscript. All authors edited and commented on the manuscript.

## METHODS

### Key Resources Table

| REAGENT or RESOURCE | SOURCE | IDENTIFIER |
| --- | --- | --- |
| **Antibodies** |  |  |
| Goat anti-human IgG-alkaline phosphatase (AP) | Southern Biotech | Cat# 2040-04 |
| Human CD19 MicroBeads | Miltenyi Biotec | Cat# 130-050-301 |
| Anti-flag-APC | BioLegend | Cat# 637307 |
| Anti-flag-PE | BioLegend | Cat# 637310 |
| Anti-His-PE | BioLegend | Cat# 362603 |
| Anti-human-IgG-PerCP-Cy5.5 | BioLegend | Cat# 410710 |
| Anti-human-CD27-APC-Cy7 | BioLegend | Cat# 356424 |
| Anti-human-CD19-BV510 | BioLegend | Cat# 302242 |
| Anti-human-IgD-FITC | BioLegend | Cat# 348206 |
| Anti-human-IgM-BV605 | BioLegend | Cat# 314524 |
| Anti-human-IgG-Alexa Fluor 647 | Thermo Fisher Scientific | Cat# A-21445 |
| HRP-conjugated mouse anti-human IgG secondary antibody | Sigma-Aldrich | Cat# RABHRP3 |
| mAb 1C7C7 | Mount Sinai, Carreno *et al.*, 2022<sup>38</sup> | ZMS1075 |
| mAb CR9114 | Dreyfus *et al.*, 2012<sup>39</sup> | N/A |
| **Bacterial and virus strains** |  |  |
| SARS-CoV-2 WA1/2020 virus | BEI Resources | NR-52281 |
| SARS-CoV-2 XBB.1.5 (hCoV-19/USA/ NY-MSHSPSP-PV76648/2022) | PVI, Mount Sinai | PV76648 |
| SARS-CoV-2 JN.1 (hCoV-19/USA/NYMSHSPSP-PV96109/2023) | PVI, Mount Sinai | PV96109 |
| **Biological samples** |  |  |
| Plasmablasts from XBB.1.5 mRNA and protein-based vaccinated donors | Fantin *et al.*, 2024<sup>20</sup> | N/A |
| Plasmablasts and memory B cells from SARS-CoV-2 mRNA vaccinated donors | Wesemann laboratory | N/A |
| Serum from SARS-CoV-2 vaccinated and convalescent donors (Cohort-3) | Simon laboratory | N/A |
| **Chemicals, peptides, and recombinant proteins** |  |  |
| SARS-CoV-2 WA1/2020 spike protein | Krammer laboratory | N/A |
| 229E HCoV spike protein | Krammer laboratory | N/A |
| HKU1 HCoV spike protein | Krammer laboratory | N/A |
| OC43 HCoV spike protein | Krammer laboratory | N/A |
| NL63 HCoV spike protein | Krammer laboratory | N/A |
| MERS-CoV spike protein | Krammer laboratory | N/A |
| SARS-CoV-2 spike protein | Krammer laboratory | N/A |
| SARS-CoV-2 WA1/2020 S2 protein | Bajic laboratory | N/A |
| SARS-CoV-2 WA1/2020 RBD protein | Krammer laboratory | N/A |
| SARS-CoV-2 WA1/2020 NTD protein | Bajic laboratory | N/A |
| SARS-CoV-2 spike protein (sorting) | Genscript | Cat# Z03481 |
| 10x phosphate buffered saline (PBS) | Boston BioProducts | Cat# BM-220-10XS |
| Glycine | Sigma-Aldrich | Cat# G7126 |
| Zinc chloride | Sigma-Aldrich | Cat# 793523 |
| Magnesium chloride | Sigma-Aldrich | Cat# M0250 |
| Tween-20 | Sigma-Aldrich | Cat# P1379 |
| Ficoll-Paque PLUS | Cytiva | Cat# 17144003 |
| 4',6-diamidino-2-phenylindole, dihydrochloride (DAPI) | Thermo Fisher Scientific | Cat# D1306 |
| Dithiothreitol (DTT) | Thermo Fisher Scientific | Cat# R0861 |
| RNaseOUT | Thermo Fisher Scientific | Cat# 10777-019 |
| Random hexamer primer | Thermo Fisher Scientific | Cat# FERSO142 |
| 10 mM dNTPs | Promega | Cat# U1515 |
| IGEPAL CA-630 | Sigma-Aldrich | Cat# I8896 |
| SuperScript III reverse transcriptase | Thermo Fisher Scientific | Cat# 18080085 |
| HotStarTaq DNA polymerase | QIAGEN | Cat# 203205 |
| Protein A agarose resin | Gold Biotechnology | Cat# P-400-5 |
| o-phenylenediamine dihydrochloride (OPD) | Sigma-Aldrich | Cat# P4664 |
| Phosphate Citrate Buffer (PCB) | Sigma-Aldrich | Cat# P4809 |
| Hydrochloric Acid | Fisher | Cat# S25856 |
| HRP-conjugated streptavidin | Pierce | Cat# 21130 |
| Dulbecco's modified eagle medium (DMEM)<sup>TM</sup> | Gibco | Cat# 11995065 |
| Minimum essential medium (MEM) amino acids solution | Gibco | Cat# 11130051 |
| Penicillin-streptomycin | Gibco | Cat# 15140122 |
| Puromycin | Invivogen | Cat# ant-pr-1 |
| Normocin | Invivogen | Cat# ant-nr-5 |
| 10% Formaldehyde | Polysciences | Cat# 04018-1 |
| Water for injection (WFI) | Gibco | Cat# A12873-01 |
| Fluorinated octyl maltoside | Anatrace | Cat# O310F |
| **Critical commercial assays** |  |  |
| ExpiFectamine293<sup>TM</sup> transfection kits | Gibco | Cat# A14525 |
| Pierce<sup>TM</sup> Antibody Biotinylation Kit | Pierce | Cat# 90407 |
| **Deposited data** |  |  |
| Paired heavy/light chain sequences (Cohort 1) | Fantin *et al.*, 2024<sup>20</sup> | BioProject: PRJNA1134144 |
| Paired heavy/light chain sequences (Cohort 2) | This paper | N/A |
| Electron microscopy maps of M15 in complex with S2 | This paper | Accession EMD-48507 |
| CryoEM structure of M15 Fab in complex with S2 | This paper | PDB ID: 9MPW |
| **Experimental models: Cell lines** |  |  |
| Expi293<sup>TM</sup> | Thermo Fisher | Cat# 14527 |
| Vero.E6.TMPRSS2<sup>TM</sup> | BPS Biosciences | Cat # 78081 |
| **Experimental models: Organisms/strains** |  |  |
| B6.Cg-Tg(K18-ACE2)2Primn/J mice | The Jackson Laboratory | Cat# 034860 |
| **Oligonucleotides** |  |  |
| Primer set for Ig PCR (Tiller *et al.*, 2008) | IDT DNA | N/A |
| **Recombinant DNA** |  |  |
| Plasmid pTwist mammalian expression vectors | Twist Biosciences | https://www.twistbioscience.com/ |
| **Software and algorithms** |  |  |
| IgBLAST | Ye *et al.*, 2011<sup>40</sup> | https://www.ncbi.nlm.nih.gov/igblast/ |
| Cell Ranger (v7.1.0) | 10X Genomics | https://www.10xgenomics.com/ |
| ImmCantation | Gupta *et al.*, 2015<sup>41</sup> | https://immcantation.readthedocs.io/en/stable/index.html# |
| CoV-AbDAb | Raybould *et al.*, 2021<sup>42</sup> | https://opig.stats.ox.ac.uk/webapps/covabdad/ |
| Observed Antibody Space (OAS) | Olsen *et al.*, 2022<sup>24</sup> | https://opig.stats.ox.ac.uk/webapps/oas/ |
| COBALT Multiple Sequence Alignment tool | Papadopoulos *et al.*, 2007<sup>43</sup> | https://www.ncbi.nlm.nih.gov/tools/cobalt/re_cobalt.cgi |
| RStudio (v4.2.1) | RStudio | https://posit.co/download/rstudio-desktop/ |
| Perl | Programming Perl | https://www.perl.org/ |
| GraphPad Prism (v10.2.3) | GraphPad | https://www.graphpad.com/ |
| MotionCorr2 | Zheng *et al.*, 2017<sup>44</sup> | N/A |
| CryoSPARC (v3.3.1) | Structura Biotechnology | https://cryosparc.com/ |
| DeepEMhancer | Sanchez-Garcia *et al.*, 2021<sup>45</sup> | N/A |
| ModelAngelo | Jamali *et al.*, 2024<sup>46</sup> | N/A |
| COOT | Emsley *et al.*, 2010<sup>47</sup> | N/A |
| Privateer | Agirre *et al.*, 2015<sup>48</sup> | N/A |
| Phenix | Leibschner *et al.*, 2015<sup>49</sup> | N/A |
| MolProbity | Williams *et al.*, 2018<sup>50</sup> | N/A |
| SBGrid | SBGrid consortium | https://sbgrid.org/ |
| Cloanalyst | Kepler *et al.*, 2013<sup>51</sup> | N/A |
| **Others** |  |  |
| Anti-human IgG Fc biosensors | Sartorius | Cat# 18-5060 |
| UltrAuFoil gold R1.2/1.3 grids | Quantifoil | Cat# TEM-Q350AR13A |

### Human samples

#### Cohort 1

Samples were collected from three individuals who received either Pfizer’s Comirnaty or Moderna’s Spikevax XBB.1.5 mRNA-based vaccine and two individuals who received Novavax’s protein-based XBB.1.5 vaccine as described in our previous study^20^. Plasmablasts were sorted 6-7 days post-vaccination as detailed in Fantin *et al.*, 2024.

#### Cohort 2

Blood samples were collected from COVID-19 infection naive individuals immunized with the COVID-19 mRNA vaccine (n=19). A longitudinal study was conducted where blood was collected one month after the second dose, six months after second dose, and 10 days after the third dose. The vaccination type, vaccination dates and history of COVID-19 were collected from each participant. All procedures involving human samples were approved by the Massachusetts General Brigham (MGH) Institutional Review Board (Protocol #:2020P000837). Informed consent was collected from participants.

#### Cohort 3

The human subjects’ samples used in this paper are sourced from two IRB approved observational research study protocols (STUDY-16-01215/IRB-16-00971 and STUDY-20-00442 /IRB-20-03374) that collect samples before and after viral antigen exposure. All study participants provided informed consent for participation in research prior to data or sample collection. All human subjects research is reviewed and approved by the Program for the Protection of Human Subjects at the Icahn School of Medicine at Mount Sinai. For set up and optimizations of the experimental work, additional samples were used from enrolled participants who provided samples prior to the SARS-CoV-2 pandemic.

Clinical metadata annotating the biospecimen at the time of collection was compiled through self-reported data from participants. Sera collected under the IRB approved protocols listed above were leveraged in this analysis. 24 sera samples from participants prior to SARS-CoV-2 vaccine approval (SARS-CoV-2 antibody negative), 25 sera samples from participants prior to SARS-CoV-2 vaccination but with evidence of a SARS-CoV-2 infection, 24 sera samples from participants who had completed primary SARS-CoV-2 immunization (2 doses of the mRNA SARS-CoV-2 vaccine) and 12 sera samples from participants who received two additional vaccinations after primary immunization were collected. Demographics, COVID-19 vaccination and SARS-CoV-2 infection information are summarized in **Table S1**.

### Plasma and peripheral blood mononuclear cell (PBMC) isolation

Blood samples from Cohort 2 participants was collected in EDTA tubes. Blood was centrifuged at 300 g for 10 minutes. Plasma was collected from the top layer and further centrifuged at 1000 g for 10 minutes for the removal of debris. Plasma was divided into aliquots and stored in -80°C. PBMCs were collected from the lower layer of the initial centrifuge step. Ficoll-Paque (Cytiva) was added to the lower layer for PBMC separation at 2200 rpm for 20 minutes. PBMCs were washed with PBS and resuspended in FBS containing 10% dimethyl sulfoxide (DMSO). Samples were aliquoted and stored at -80°C and liquid nitrogen.

### Flow Cytometry and single-cell sorting

PBMCs stored at -80°C were thawed at 37°C and transferred to warm RPMI 1640 media (Gibco) supplemented with 10% FBS (Cytiva). B cells were enriched via positive selection of with anti-CD19 beads (Miltenyi). As previously described^52^, spike-positive memory B cells were sorted (BD FACSAria Fusion) into 96-well plates. In brief, enriched B cells were stained with SARS-CoV-2 spike containing a Flag tag (Genscript). Following primary incubation, the enriched B cells were stained with allophycocyanin (APC)-conjugated anti-Flag and phycoerythrin (PE)-conjugated anti-Flag for double positive sorting. DAPI^-^ CD20^+^ CD19^+^ IgM^-^ IgD^-^ IgG^+^ CD27^+^ Spike^+^ cells for memory B cell, and DAPI^-^ CD20^-^ CD19^+^ CD38^hi^ CD27^+^ cells for plasmablasts were single cell sorted into 96-well PCR plates containing lysis buffer (4 μL of 0.5x PBS with 10 mM dithiothreitol and 4U of RNAseOUT). Plates were stored at -80°C.

### Bioinformatic analysis of B cell receptor (BCR) sequencing data and database mapping

Heavy and light chain single-cell V(D)J sequences were obtained from plasmablasts from Cohort 1 after running 10X Genomics Cell Ranger v7.1.0 on FASTQ files generated via single-cell RNA (sc-RNA) sequencing, as described by Fantin and colleagues^20^. For Cohort 2, as previously described^52^, RNA from single cell sorted spike positive memory B cells were reverse transcribed into cDNA for amplification of IgH, Igκ, and Igƛ genes. In brief, cDNA was amplified using two semi-nested PCRs^53^. Sanger sequencing was performed on the PCR products from the second amplification. Cells with unique and productive heavy and light chains were used for further analysis. Each paired sequence was annotated with its V(D)J genes using the AssignGenes and ParseDb functions in the changeo module of the Immcantation framework^41^. Paired VH/VL sequences from the CoV-AbDab and OAS databases were downloaded from the version updates on November 14, 2023, and June 21, 2024, respectively^24,42^. For CoV-AbDab, sequences with undefined V or J genes or CDR3 sequences for heavy and light chains and those from species other than humans were filtered out. Sequence-based mapping to databases was performed in R, as follows: a sequence from the cohort was mapped to a sequence in the database if i) they had the same V and J gene annotations in both heavy and light chains and ii) they had a CDR3 amino acid similarity greater than or equal to 70% in both heavy and light chains^23^. The same matching criteria were followed while mapping M15 from Cohort 1 to Cohort 2.

### Sequence analysis

M15-like VH and VL amino acid sequences were aligned using the COBALT multiple alignment tool (https://www.ncbi.nlm.nih.gov/tools/cobalt/re_cobalt.cgi). Mutation frequencies within clonal lineages and other M15-like sequences were assessed from FASTA-formatted files using R and Perl.

### Monoclonal antibody production and purification

The M15 mAb was produced based on the protocol described by Fantin and colleagues^20^. Briefly, the VH and VK genes of M15 were cloned into pTwist mammalian expression vectors. Expi293 cells (25-to 200-mL cultures) were transfected with the heavy- and light-chain-expressing vectors at a 1:2 ratio by concentration, along with Expifectamine (transfection reagent; Thermo Fisher Scientific), following the protocol described in the lipid-based Thermo Fisher Expi293™ Expression System Kit. The cells were incubated in a shaker flask at 37°C for 18-22 hours at 125 rpm. Enhancers 1 and 2 were added to the culture the next day, followed by further incubation for 5-6 days. Cells were then pelleted down by centrifugation and filtered (0.45 µm). The supernatant was mixed with protein A agarose resin (Gold Biotechnology) in a rotary shaker for 12 hours at 4°C and then loaded onto Poly-Prep chromatography columns (Bio-Rad Laboratories, Inc.) and eluted with glycine in phosphate-buffered saline (PBS), followed by neutralization with 1 M Tris pH 8 and 5 M NaCl. Purified antibodies were concentrated and stored at 4°C.

### ELISA

For ELISA, 96-well Immulon 4 HBX plates (Thermo Scientific) were coated with 50 µL of the recombinant protein at 2 µg/mL and were stored at 4°C overnight. Wells were then washed three times with phosphate-buffered saline (3) containing 0.1% Tween-20 (PBST) and blocked with 100 µL PBST supplemented with 3% milk for 1 hour at room temperature. Antibodies were diluted 3-fold from a starting concentration of 30 µg/mL, and sera were diluted 2-fold from a starting concentration of 1:25, all in PBST supplemented with 1% milk in a separate 96-well round-bottomed plate. For the vaccination (4X) sera samples, a starting dilution of 1:50 was used. After the blocking solution was discarded, 100 µL of the diluted antibodies or sera was added to the corresponding wells, followed by incubation for 1 hour (antibodies) or 2 hours (sera) at room temperature. The antibody or sera solutions were then discarded, and the wells were washed three times with PBST. Next, 100 µL of mouse anti-human IgG antibody conjugated to horseradish peroxidase (Sigma, A0293) at a 1:3000 dilution was added to each well, followed by incubation for 1 hour at room temperature. Plates were washed three times with PBST and developed by adding 100 µL of o-phenylenediamine dihydrochloride (OPD; Sigma-Aldrich) in phosphate citrate buffer (PCB; Sigma-Aldrich) with 0.04% hydrogen peroxide for 10 minutes. The reaction was stopped by adding 50 µL of 3 M hydrochloric acid (HCl; Fisher). The optical density (OD) was read for each well using a Synergy 4 (BioTek) plate reader at a wavelength of 490 nm. AUC values were computed in GraphPad Prism (v10.2.3), setting the average plus five times the standard deviation of the OD of blank wells as the baseline.

### Biolayer interferometry (BLI)

Anti-human IgG Fc biosensors (Sartorius; 18-5001) were hydrated in 10 nM Tris, 150 mM NaCl, and 0.1% Tween buffer for 10 minutes. Then, 2 µg/mL M15 diluted in the same buffer was immobilized on a biosensor, and purified recombinant S2 protein was flowed at different concentrations, starting from 166.7 nM to 16,666.7 nM, using the ForteBio Sartorius BLItz System. The steps involved are briefly described as follows: i) Baseline: the biosensor was initialized at baseline by dipping it in buffer for 30 seconds, ii) Loading: 4 µL of the antibody (M15) at 2 µg/mL was loaded onto the biosensor for 90 seconds, iii) Baseline: the biosensor was dipped back in buffer for 30 seconds, iv) Association: 4 µL of the antigen (S2) at different concentrations was loaded onto the biosensor for 60 seconds, and v) Dissociation: the biosensor was dipped back into the buffer to let the antigen dissociate for 90 seconds. A control run with buffer loaded instead of the antibody was performed to confirm specific binding. The R_max-apparent_ values for association, with the baseline signal before association subtracted, were plotted against the S2 concentrations, and the K_D_ was determined using a non-linear fit for one-site specific binding in GraphPad Prism (v10.2.3).

### Linkage analysis

For each clonal lineage, association or linkage between each pair of convergent mutations was assessed by constructing a contingency table of co-occurrence of the two mutations. The O.R. was computed by running the fisher.test() function on the contingency table using the R package ‘stats’.

### Serum competition ELISA

For serum competition ELISA, 96-well Immulon 4 HBX plates (Thermo Scientific) were coated with 50 µL of the recombinant spike protein at 2 µg/mL and stored at 4°C overnight. Wells were then washed three times with PBST and blocked with PBST supplemented with 3% milk for 1 hour at room temperature. Serum was diluted 2-fold from a starting dilution of 1:25 in PBST supplemented with 1% milk in a separate 96-well round-bottomed plate. After the blocking solution was discarded, 100 µL of the diluted serum was added to the corresponding wells, followed by incubation for 2 hours at room temperature. For the mAb-only positive control, PBST supplemented with 1% milk was added instead of serum. After incubation, the serum solutions or control solution was discarded, and the wells were washed three times with PBST. For each serum sample, 100 µL of biotinylated M15 antibody, or an mpox-specific antibody as a negative control, was added to each dilution in duplicates at two times the minimum binding concentration of M15 with spike protein (quantified by ELISA), followed by incubation for 2 hours at room temperature. The antibody solutions were then discarded, and the wells were washed three times with PBST. Next, 100 µL of streptavidin conjugated to horseradish peroxidase (Pierce, 21130) at a 1:1000 dilution was added to each well, followed by incubation for 1 hour at room temperature. Plates were washed three times with PBST and developed by adding 100 µL of OPD (Sigma-Aldrich) in PCB (Sigma-Aldrich) with 0.04% hydrogen peroxide for 10 minutes. The reaction was stopped by adding 50 µL of 3 M HCl (Fisher). The absorbance was read for each well using a Synergy 4 (BioTek) plate reader at a wavelength of 490 nm. Competition ED_50_ values were computed by fitting the OD-dilution curves with the ‘Absolute IC_50_’ function in GraphPad Prism (v10.2.3).

### Neutralization assay

Vero E6 cells expressing transmembrane protease serine 2 (TMPRSS2; BPS Biosciences, 78081) were maintained in Dulbecco’s modified Eagle medium (DMEM; Gibco) supplemented with 10% fetal bovine serum (FBS), 1% minimum essential medium (MEM) amino acids solution (Gibco, 11130051), 100 units/mL penicillin and 100 µg/mL streptomycin (Gibco, 15140122), 3 µg/mL puromycin (InvivoGen, ant-pr-1), and 100 µg/mL normocin (InvivoGen, ant-nr-05). Viral stocks were confirmed through sequencing and quantified using the 50% tissue culture infectious dose (TCID_50_) method. Microneutralization assays were performed as previously detailed^38^. In brief, Vero E6 TMPRSS2 cells were seeded in 96-well tissue culture plates (Corning, 3340) at a density of 1 × 10^4^ cells per well. After 24 hours, mAbs were diluted to an initial concentration of 30 µg/mL in 1x MEM with 2% FBS and then serially diluted 1:3 to achieve a final concentration of 0.041 µg/mL. An 80-µL portion of the mAb dilution was mixed with 80 µL of SARS-CoV-2 diluted to 10^4^ TCID_50_/mL, and this mixture was incubated at room temperature for 1 hour. Following incubation, 120 μL of the virus/mAb mixture was used to infect the cells for 1 hour. The inoculum was subsequently removed and replaced with 100 µL of 1x MEM containing 2% FBS and 100 µL of antibody dilution. The cells were then incubated at 37°C in a 5% CO_2_ environment for 48 hours before fixation with 10% paraformaldehyde (Polysciences) for 24 hours. After fixation, the paraformaldehyde was discarded, and cells were permeabilized by adding 100 µL of PBS containing 0.1% Triton X-100 (Fisher) for 15 minutes at room temperature. Once permeabilization was complete, the Triton X-100 solution was removed, and cells were blocked with 100 µL of 3% non-fat milk (Life Technologies) diluted in PBS. Finally, cells were stained for SARS-CoV-2 nucleoprotein using mAb 17C7, as previously described^38^.

### Mice protection studies

Animal studies were performed in accordance with the guidelines set forth by the Icahn School of Medicine at Mount Sinai’s Institutional Animal Care and Use Committee (IACUC) and under approved protocols reviewed by the IACUC. The 50% lethal dose (LD_50_) was determined for WA1/2020, XBB.1.5, and JN.1 by infecting 6- to 8-week-old hACE2-K18 mice (Jackson Laboratory) with serial dilutions of the virus, ranging from 10^5^ to 5 plaque-forming units (PFU) in sterile PBS. For the protection studies, mice received an intraperitoneal injection of mAb at a dosage of 10 mg/kg, diluted in 100 µL of sterile PBS. Two hours post-treatment, mice were anesthetized using 0.15 mg/kg ketamine and 0.03 mg/kg xylazine, both diluted in water for injection (WFI, Gibco), and intranasally infected with a 3xLD_50_ dose of SARS-CoV-2 WA1/2020, XBB.1.5, or JN.1 diluted in 50 µL of sterile PBS. Mice were monitored for weight loss for 14 days following infection, and any animals that lost more than 25% of their body weight were humanely euthanized. The influenza A virus antibody CR9114 served as a negative control^39^.

### Cryogenic electron microscopy (Cryo-EM) sample preparation

SARS-CoV-2 stabilized S2 domain was incubated with M15 Fab at 2.5 mg/mL at a molar ratio of 1.5:1 Fab:S2 for 20 minutes at 4°C. Immediately before grid preparation, fluorinated octyl maltoside was added to the pre-formed complex at 0.02% w/v final concentration. Then, 3-µL aliquots were applied to UltrAuFoil gold R1.2/1.3 grids (Quantifoil) and subsequently blotted for 6 seconds at blot force 1 and then plunge-frozen in liquid ethane using an FEI Vitrobot Mark IV. Grids were imaged on a Titan Krios microscope operated at 300 kV and equipped with a Gatan K3 Summit direct detector. 8,134 movies were collected in counting mode at 16e−/pix/s at a magnification of 105,000, corresponding to a calibrated pixel size of 0.826 Å. Defocus values were from -0.5 to -1.5 µm.

### Cryo-EM data processing

Movies were aligned and dose-weighted using MotionCorr2^44^. Contrast transfer function estimation was done in cryoSPARC v3.3.1 using Patch CTF, and particles were picked with cryoSPARC’s blob picker. The picked particles were extracted with a box size of 512 pixels, with 4x binning, and subjected to a 2D classification. Selected particles were then subjected to a second round of 2D classification. An initial model was generated on 164,341 selected particles at 6 Å/pixel with 4 classes. The best class, containing 74,661 particles, was selected for further processing. After one round of non-uniform refinement, without imposed symmetry, the particles were subjected to 3D classification with 5 classes. Of these, the best 3 classes, containing 321,918 particles in total, were combined, re-extracted without binning with a box size of 512 pixels, and selected for further rounds of non-uniform refinement with local CTF refinement, yielding the final global map at a nominal resolution of 2.53 Å. The protomer with the best Fab volume was subjected to local refinement with a soft mask extended by 6 pixels and padded by 12 pixels encompassing S2 and Fab. A second round of local refinement was performed with a soft mask encompassing the receptor-binding domain and variable domains of the Fab. This yielded the final local map at 3.16 Å resolution. The two half-maps from the global or local refinement were used for sharpening in DeepEMhancer^45^. The reported resolutions are based on the gold-standard Fourier shell correlation of 0.143 criterion.

### Atomic model building and refinement

The DeepEMhancer sharpened map was used for model building with ModelAngelo^46^ and then manually built using COOT^47^. N-linked glycans were built manually in COOT using the glyco extension and their stereochemistry and fit to the map validated with Privateer^48^. The model was then refined in Phenix^49^ using real-space refinement and validated with MolProbity^50^. The structural biology software was compiled and made available through SBGrid^54^.

### Clonal lineage inference

Clonal lineages using paired heavy and light chain M15-like sequences from OAS and CoV-AbDab were constructed, and the UCA and intermediate sequences were inferred using Cloanalyst^51^.

## SUPPLEMENTARY MATERIAL

**Supplementary Table 1.** Demographic information for Cohorts 1, 2 (sequences of human antibodies post COVID-19 vaccination) and 3 (human sera samples used in competition ELISA).

| Demographic | Cohort 1<br>n = 5 | Cohort 2<br>n = 19 | Cohort 3<br>n = 85 |
| --- | --- | --- | --- |
| **Age groups (number of individuals)** |  |  |  |
| 18-29 | 0 | 3 | 21 |
| 30-39 | 1 | 9 | 23 |
| 40-49 | 2 | 4 | 18 |
| 50-59 | 1 | 1 | 14 |
| 60-69 | 1 | 1 | 4 |
| 70-79 | 0 | 1 | 5 |
| **Sex at birth** | 3M, 2F | 6M, 13F | 33M, 52F |
| **Previous immune history to SARS-CoV-2** | XBB.1.5 mRNA or protein-based vaccination, five to six previous doses of SARS-CoV-2 vaccination, none to two previous SARS-CoV-2 infection | Two or three doses of SARS-CoV-2 mRNA vaccination, no previous SARS-CoV-2 infection | Pre-pandemic (n = 24) SARS-CoV-2 convalescent (n = 25) Two doses of SARS-CoV-2 mRNA vaccination (n = 24) Four doses of SARS-CoV-2 mRNA vaccination (n = 12) |

**Supplementary Table 2.**
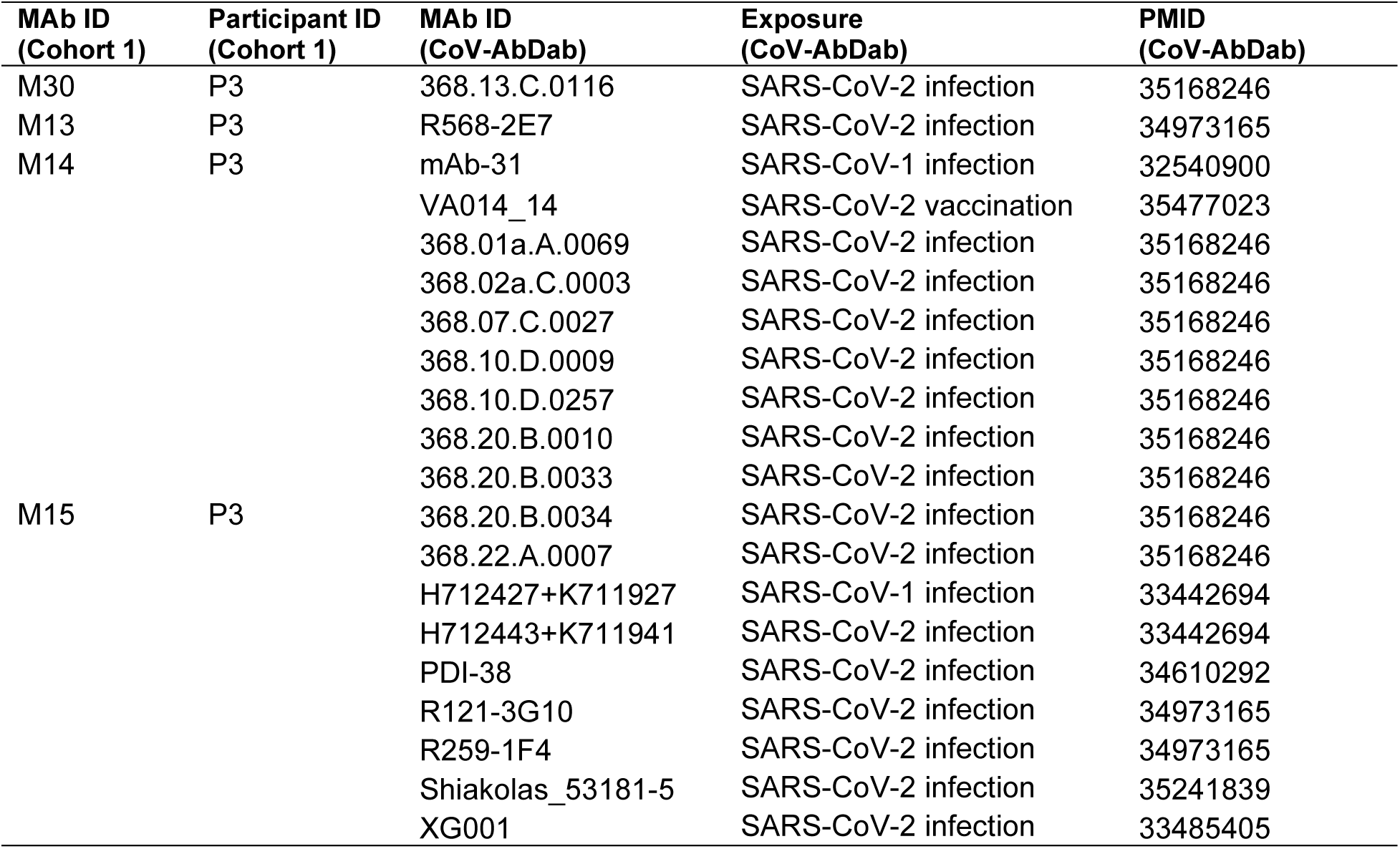
Antibodies from CoV-AbDab that match with mAb sequences from Cohort 1. Antibody sequences from Cohort 1 that mapped to mAb sequences in CoV-AbDab based on identical V and J gene usage and >70% CDR3 similarity in both heavy and light chains are shown. The exposure groups of the matched mAbs in CoV-AbDab and the references (PMIDs) for the studies from which these mAbs were retrieved in CoV-AbDab and annotated.

**Supplementary Table 3.**
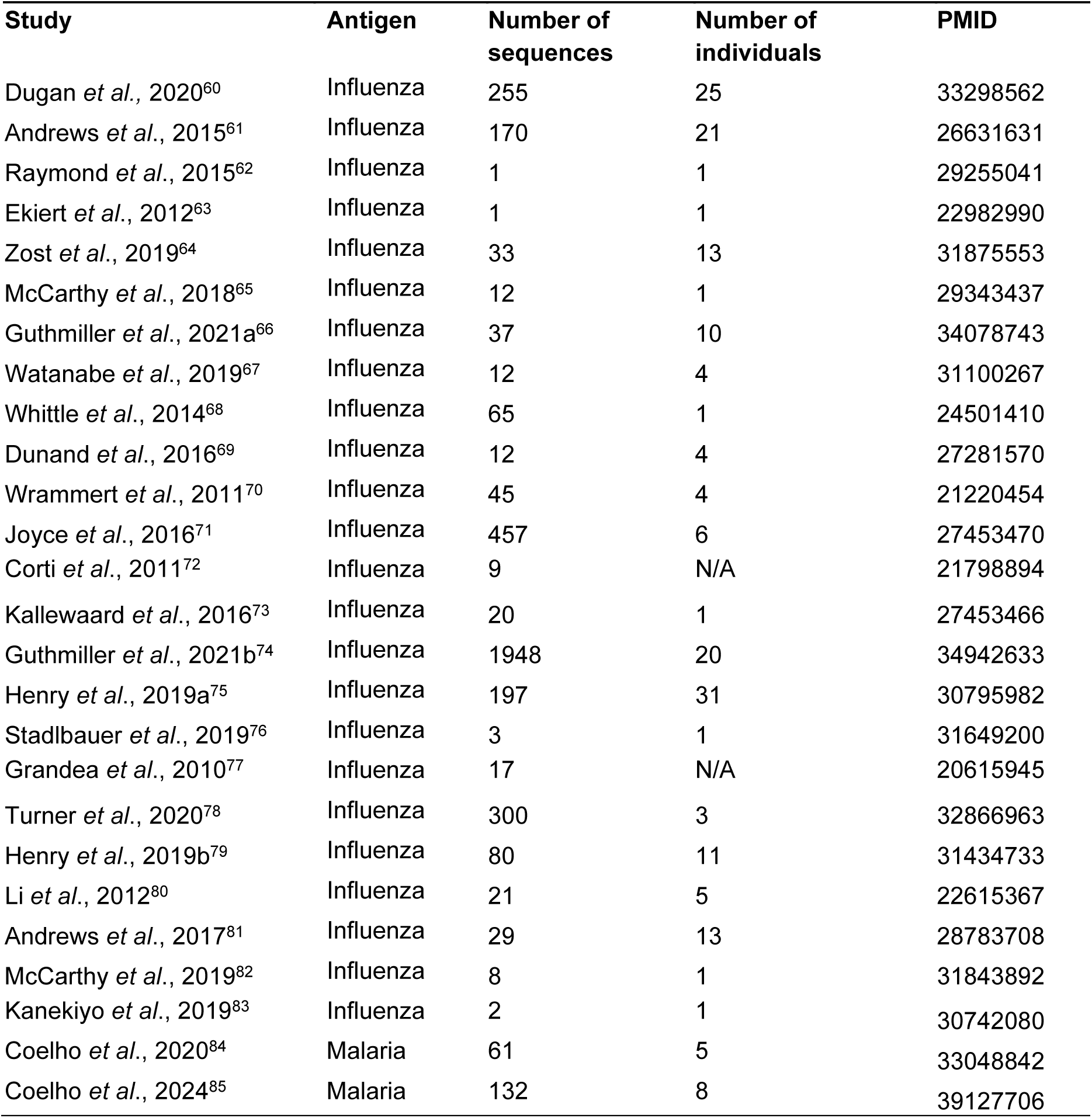
In-house compiled dataset of monoclonal antibodies (mAbs) generated in response to influenza infection, influenza vaccination, or malaria vaccination. Studies lacking information on the number of individuals have been marked as “N/A.”.

**Supplementary Figure 1.**
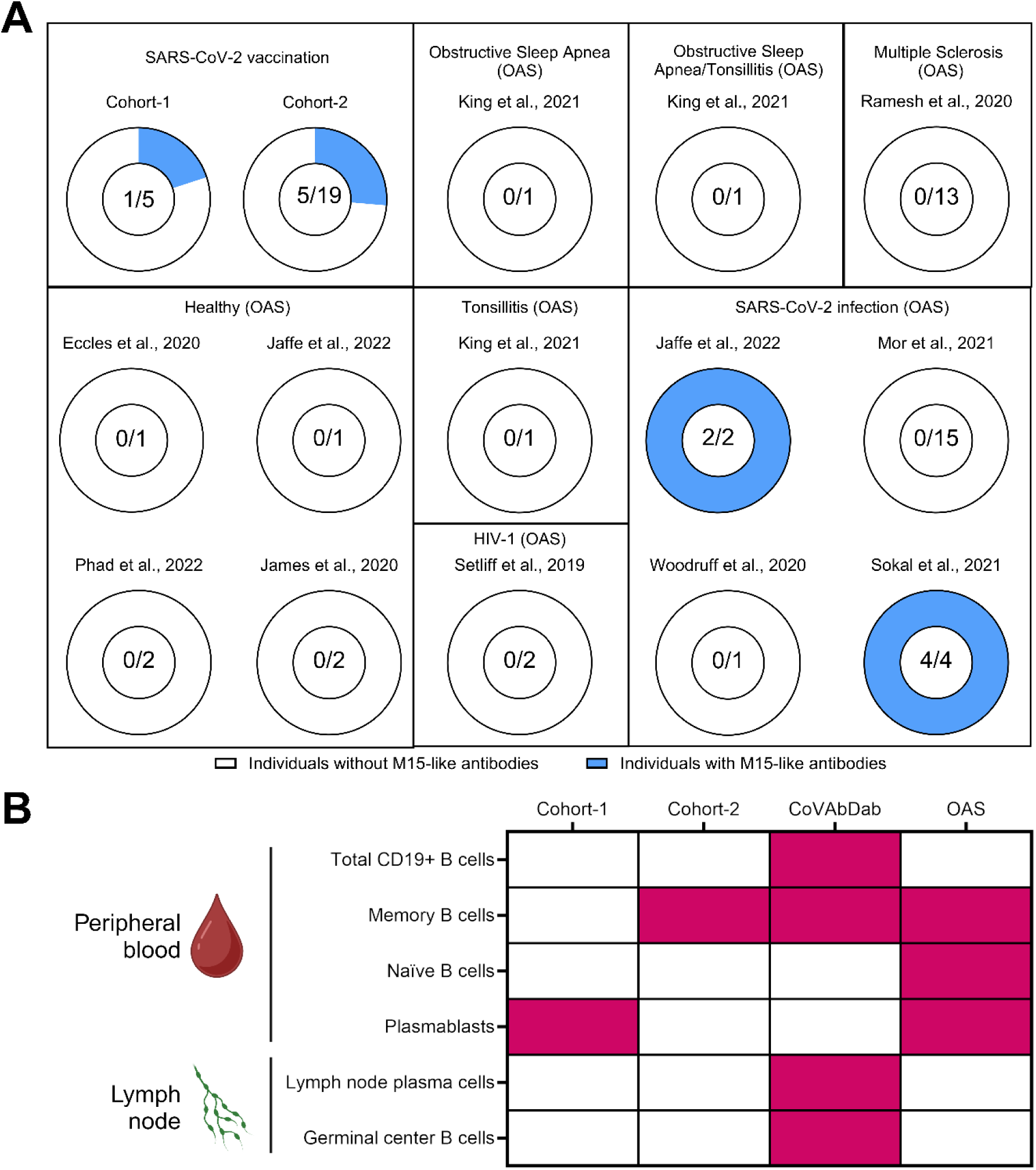
Identification of human B cells with M15-like antibody sequences. **(A)** Frequency of individuals with M15-like antibodies from Cohort 1, Cohort 2, and OAS databse, represented as donut plots. **(B)** The presence of the B cell subtype (y-axis) expressing an M15-like BCR/antibody in the corresponding cohort or database where the sequence was identified is shown in pink. Due to variability in sequencing depth across studies, a quantitative analysis comparing frequencies across anatomical sites was not feasible.

**Supplementary Figure 2.**
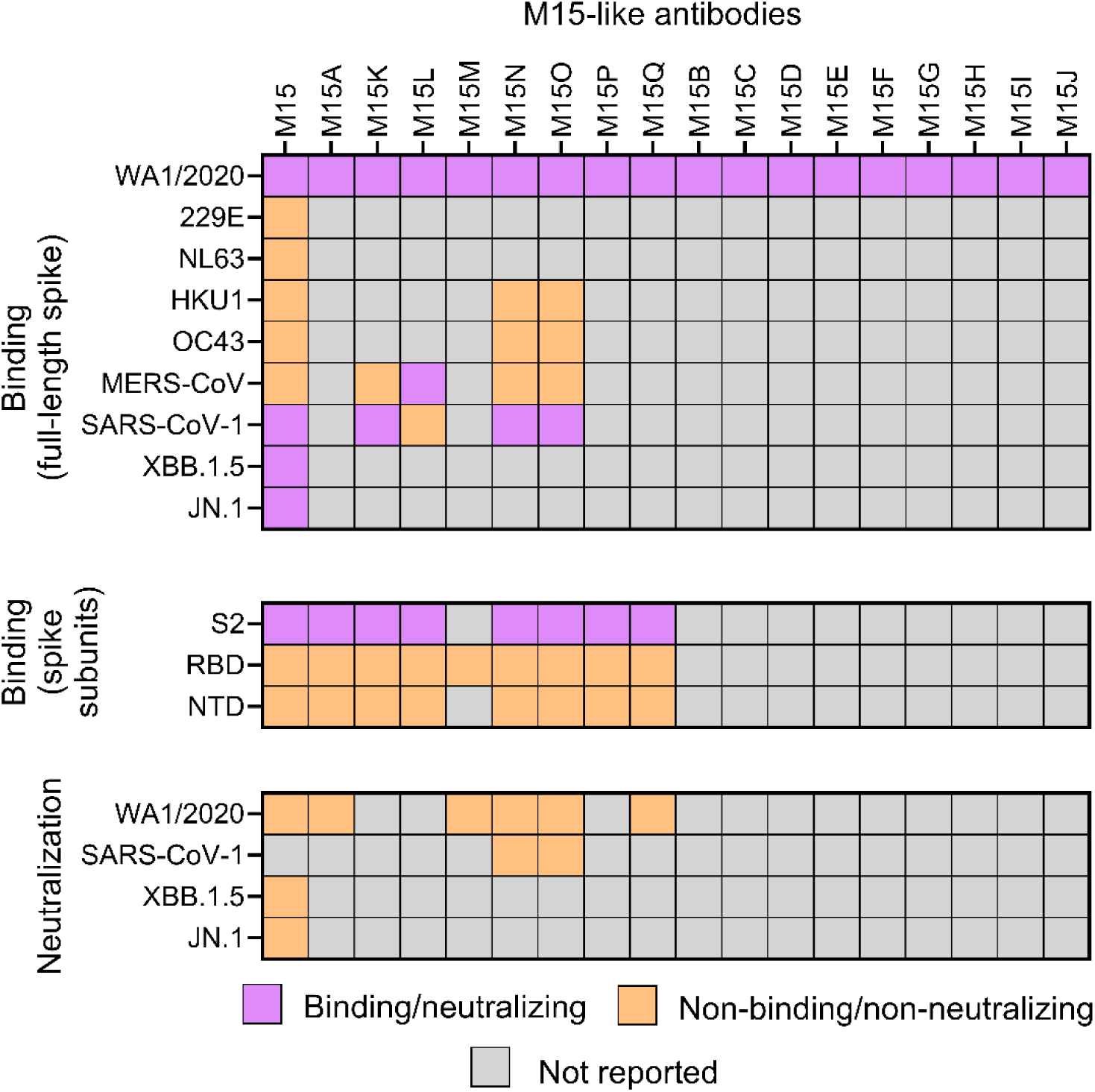
Binding and neutralization activity of M15-like antibodies from CoV-AbDab. Binding and neutralization activity of M15-like antibodies, as reported in the individual studies s in CoV-AbDab. These data confirm that M15-like antibodies are S2-targeting non-neutralizing molecules.

**Supplementary Figure 3.**
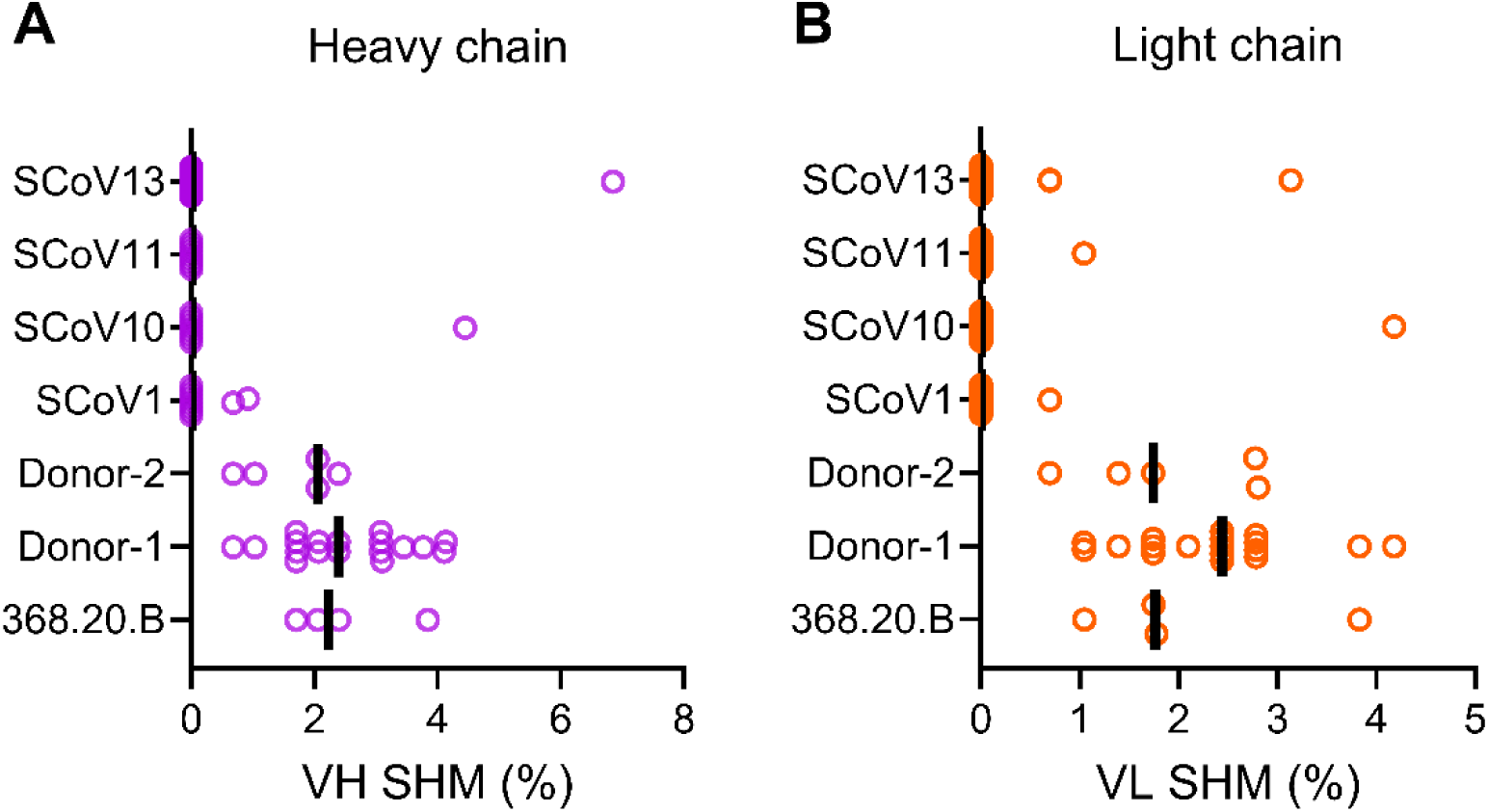
Extent of SHM in sequences from M15-like clonotypes. SHM rates in the (A) heavy and (B) light chain variable regions among the seven M15-like clonal lineages constructed from individuals with at least three M15-like sequences. The bar represents the median of SHM values.

**Supplementary Figure 4.**
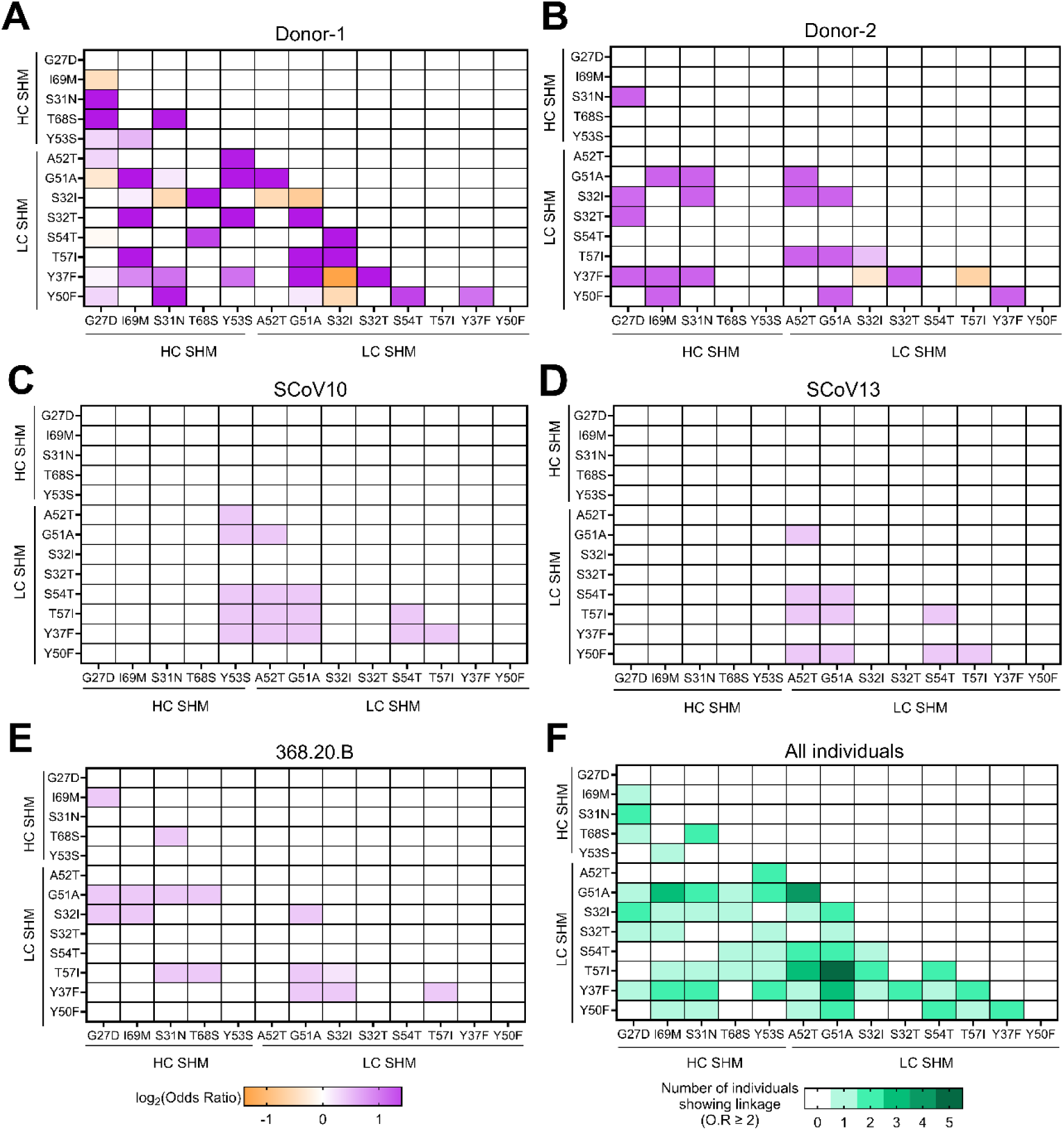
Linkage between convergent SHM in M15-like clonotypes. **(A-E)** Heatmaps representing linkage between each pair of heavy and light chain convergent mutations within each M15-like clonotype. Linkage is represented as log2 (Odds Ratio). **(F)** Heatmap representing the number of individuals showing significant linkage (defined as odds ratio (O.R) ≥2) between each pair of convergent mutations in heavy and light chains. The location of mutations (heavy/light chain) is mentioned next to or below the axes. Since the heatmap is symmetric, values in the upper right triangle are not shown. Individuals SCoV1 and SCoV11 were omitted due to lack of linkage between any pair of mutations.

**Supplementary Figure 5.**
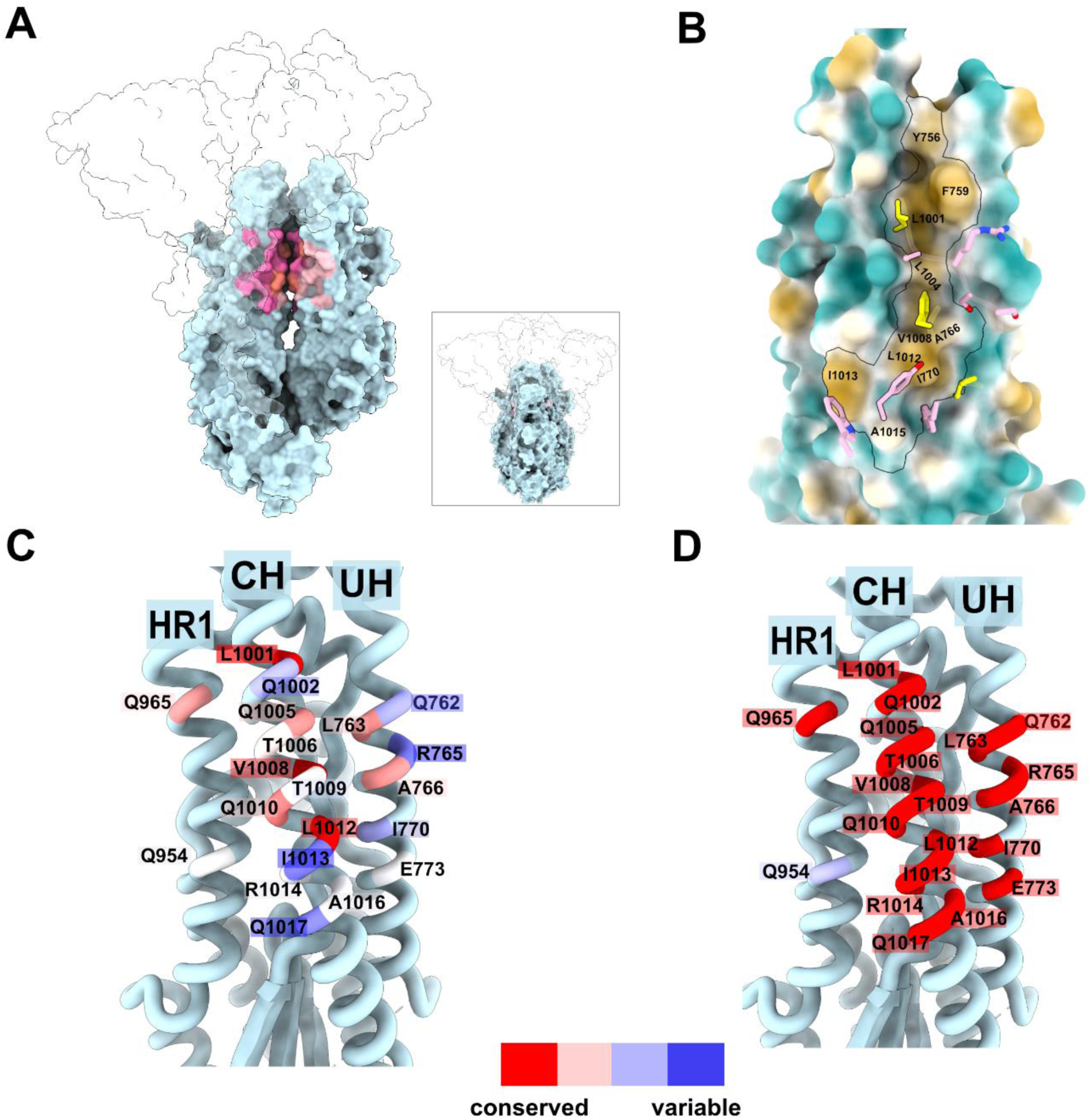
Physicochemical properties and conservation of the central interface epitope. **(A)** Position of the M15 epitope at the interprotomer interface of the full-length SARS-CoV2 spike in the canonical closed conformation (PDB 6ZP2). S1 domains and one out of three S2 protomers are rendered in a transparent white to permit visualization of the occluded M15 binding footprint. S2 residues are colored, as in figure 4C (heavy chain contacts in magenta, light chain contacts in pink). S2 Residues contacting both heavy at light chains are rendered in salmon. **(B)** S2 surface colored according to hydrophobicity (Kyte-Doolittle hydrophobicity score) of S2 residues. Hydrophobicity values are depicted in a color palette ranging from camel hair (hydrophobic) to robin’s egg blue (hydrophilic). M15 light chain contacts are displayed in pink, with convergent/specific mutations highlighted in yellow. **(C)** Conservation of M15 contacts across hCoVs 229E, NL63, HKU1, HKU4, HKU5, OC43, MERS-CoV, SARS-CoV and SARS-CoV-2, displayed on ribbon structure of S2. **(D)** Conservation of M15 contacts across sarbecoviruses SARS-CoV-1, SARS-CoV-2 WA1/2020 and SARS-CoV-2 variants B.1.1.7, P.1, B.1.617.2, BA.1, BA.2.86, BA.5, BQ.1.1, XBB.1.5 and JN.1, displayed on ribbon structure of S2.

**Supplementary Figure 6.**
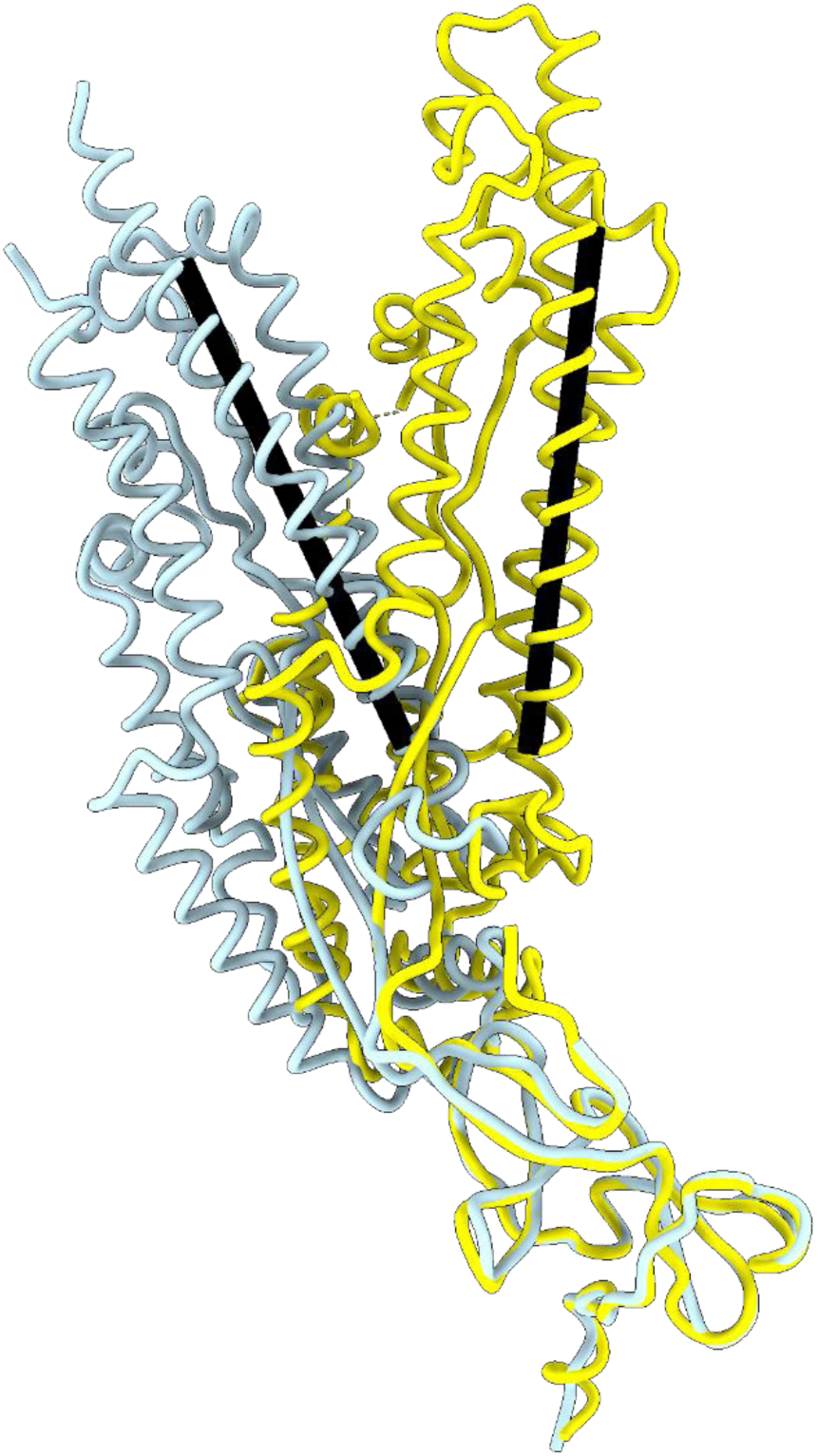
The M15-bound S2 protomer (gray) splays outwards from an axis defined by the central helix. S2 protomer structures derived from M15-S2 complex (gray) and the structure of S2 in the absence of M15 (yellow; PDB 8VQB) were superposed at residues 1010-1040. Axes were defined by the central helix between residues 998-1028 in each S2 model. The angle subtending said axes measures 32°.

**Supplementary Figure 7.**
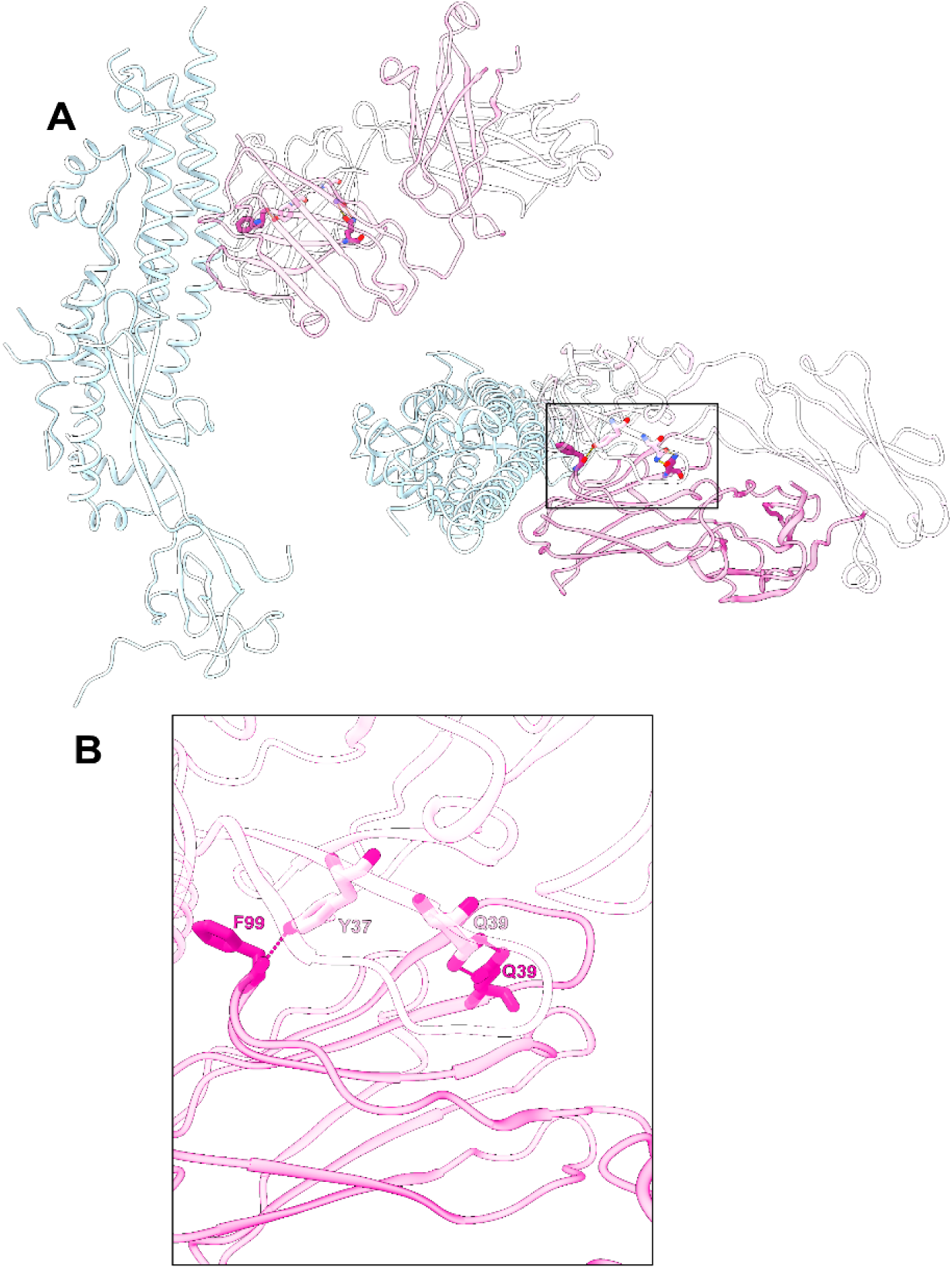
Inter-chain hydrogen bonds between residues within M15 Fab heavy and light chain variable regions. **(A)** Inter-chain contacts are displayed in stick representation, situated within the ribbon structure of M15-S2 protomer structure; lateral view, left; top-down view, right. **(B)** Close-up view of inter-Fab contacts.

**Supplementary Figure 8.**
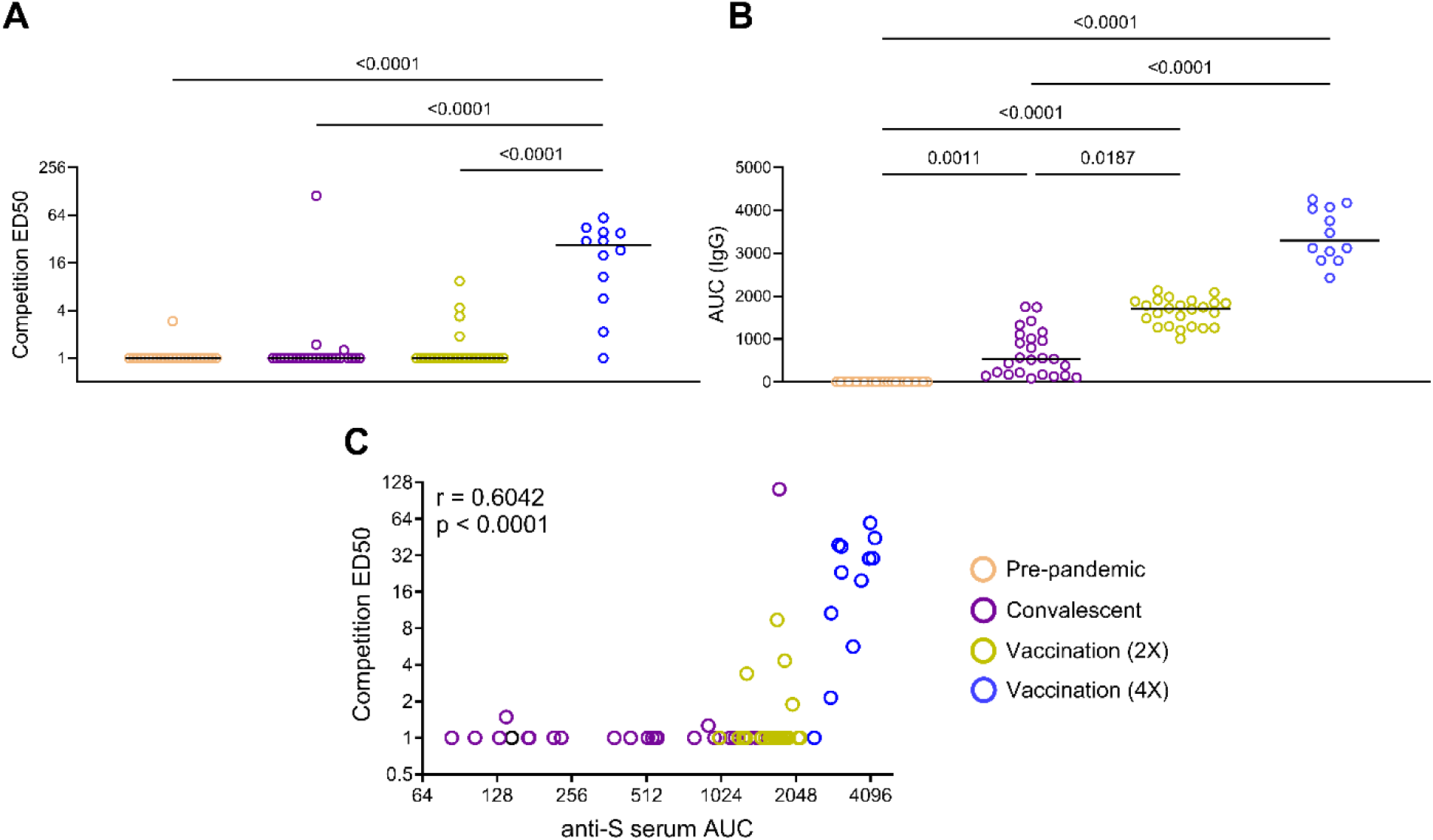
Competition ELISA between serum antibodies and M15 mAb. **(A)** Serum competition ELISA (unnormalized) between biotinylated M15 and human sera from pre-pandemic, SARS-CoV-2 convalescent, two doses of SARS-CoV-2 vaccination and four doses of SARS-CoV-2 vaccination. Competition is represented as unnormalized ED50 values. ED50 values less than 1 were set to a constant value of 1. **(B)** Anti-spike serum ELISA titers from pre-pandemic, SARS-CoV-2 convalescent, two doses of SARS-CoV-2 vaccination, and four doses of SARS-CoV-2 vaccination sera samples. Serum titers are computed as area under the curve (AUC) values. **(C)** Correlation between competition ED50 values and anti-spike serum titers computed as AUC values of all samples. The circles are colored based on exposure groups. Comparisons between exposure groups in (A) and (B) were performed using the Kruskal-Wallis test followed by Dunn’s multiple correction. Spearman correlation was performed to estimate the r-coefficient and *p*-values for the correlation analysis in (C). p-values represent total correlation across infection, two dose vaccination and four dose vaccination groups.

